# Homology Directed Repair by Cas9:Donor Co-localization in Mammalian Cells

**DOI:** 10.1101/248179

**Authors:** Philip J.R. Roche, Heidi Gytz, Faiz Hussain, Christopher J.F. Cameron, Denis Paquette, Mathieu Blanchette, Josée Dostie, Bhushan Nagar, Uri David Akavia

## Abstract

Homology directed repair (HDR) induced by site specific DNA double strand breaks (DSB) with CRISPR/Cas9 is a precision gene editing approach that occurs at low frequency in comparison to indel forming non homologous end joining (NHEJ). In order to obtain high HDR percentages in mammalian cells, we engineered Cas9 protein fused to a high-affinity monoavidin domain to deliver biotinylated donor DNA to a DSB site. In addition, we used the cationic polymer, polyethylenimine, to deliver Cas9 RNP-donor DNA complex into the cell. Combining these strategies improved HDR percentages of up to 90% in three tested loci (CXCR4, EMX1, and TLR) in standard HEK293 cells. Our approach offers a cost effective, simple and broadly applicable gene editing method, thereby expanding the CRISPR/Cas9 genome editing toolbox.

**Summary:** Precision gene editing occurs at a low percentage in mammalian cells using Cas9. Colocalization of donor with Cas9MAV and PEI delivery raises HDR occurrence.

## Introduction

CRISPR/Cas9 has revolutionized gene editing methods by offering a means to introduce DNA double strand breaks (DSBs) at specific loci. The Cas9 nuclease and a short guide RNA (sgRNA) form the Cas9/sgRNA ribonucleoprotein (RNP) complex. DSBs are repaired in the cell by non-homologous end joining (NHEJ) or by homology directed repair (HDR). NHEJ generates deletions and insertions (indels) at the target site, whereas HDR utilizes a donor DNA (dDNA) template for lesion repair. An external dDNA allows precise gene modifications such as single-point mutations (Mali *et al*, 2013; Cong *et al*, 2013) or large insertions (Schwank *et al*, 2013). Without chemical selection strategies or genetic interventions, HDR percentages below 1% are considered normal (Miyaoka *et al*, 2016). To identify cells with correctly edited genomes, current approaches select Cas9 expressing cells via fluorescent markers and/or antibiotic resistance to isolate Cas9 expressing HDR-edited single-cell clones (Mali *et al*, 2013; Cong *et al*, 2013; Schwank *et al*, 2013; Van Trung Chu *et al*, 2015). While selection raises the percentage of successfully modified clones, it requires a selection marker, and significant additional experimental time.

Inhibition of NHEJ mediators, (DNA ligase IV or Ku70/80) by small molecules (SCR7); protein-based (expression of viral Ad4); and shRNA achieve NHEJ decreases (with increases in HDR) of 4-fold, 8-fold and 5-fold, respectively ( Van Trung Chu *et al*, 2015). Cell cycle influences HDR frequency, as NHEJ occurs in G1 phase and HDR in S/G2. Nocodazole-arrested cells in the S/G2 phase exhibited 38% HDR with Cas9 RNP (Lin *et al*, 2014) but this treatment is not effective in all cell types. In the MDA-MB-4687 breast cancer cell line model for example, nocodazole, vincristine or colchicine treatment causes G1 arrest (Holt *et al*, 1997). Furthermore, while these pharmacological interventions are possible in cell culture, they can’t be used *in vivo* (Lee *et al*, 2017a; Yanik *et al*, 2017).

dDNA homology arm (HA) length and structure can influence HDR occurrence. Zhang et al. demonstrated 26% HDR using double cut plasmid donors with long (600 bp) HAs (Zhang *et al*, 2017). Cas9 complex releases the distal DNA strands (single or double stranded) almost immediately after making the cut, while the proximal strand is released later. Considering this mechanism, HDR with asymmetric ssDNA improved by 2.6-fold compared to symmetric ssDNA and by 4-fold over dsDNA (Shibata *et al*, 2017; Richardson *et al*, 2016). Thus, HA optimization increases HDR percentages up to 57%, surpassing both chemical and genetic interventions (Richardson *et al*, 2016) (further summarized in Supplementary Table 1).

The delivery method used to introduce Cas9 into a cell influences the percentage of cells modified, and can sometimes also influence the type of DNA repair between HDR and NHEJ. The current state-of-the-art is nucleofection, in which the Cas9 RNP complex is delivered directly into a cell nucleus. Nucleofection results in lower off-target DSBs (Kim *et al*, 2014; Kouranova *et al*, 2016) and reduced lag time in nuclease action (Lin *et al*, 2014), thereby increasing both NHEJ and HDR. The advantage of using a pre-assembled Cas9 RNP complex over traditional plasmid delivery is that it avoids the time lapse associated with mRNA and sgRNA transcription, Cas9 mRNA translation, and formation of intracellular RNP. Questions remain as to whether nucleofection is applicable in clinical settings other than ex-vivo editing and transplantation (Lin *et al*, 2017).

Cationic polymer RNP delivery is a flexible delivery platform that has thus far drawn limited attention in CRISPR/Cas9 applications. To date, there have been only two novel attempts using this approach in mammalian cells - sgRNA:dDNA chimeras and CRISPR-GOLD applying PAsp(DET) polyplexes (Lee *et al*, 2017b; 2017a). There has also been one recent work of using polyethylenimine (PEI) to introduce Cas9 into bacterial cells (Kang *et al*, 2017). Taking into account cost, biodegradation (Wen *et al*, 2009), and proven nucleic acid transfection utility (Boussif *et al*, 1995), Cas9 RNP delivery by PEI is a promising approach for gene editing. PEI enters the cell by size dependent entry mechanisms (clathrin, calnexin or macropinocytosis) (Khalil *et al*, 2006), escaping endosomes via a proton pump effect and translocating to the nucleus (Oh *et al*, 2002). PEI increases DNA nuclear localization over poly-L lysine polyplexes (Pollard *et al*, 1998). An additional benefit of PEI is microtubular active transport, which could increase Cas9 concentration at the peri-nucleus based on the behaviour of polyplexes in early gene transfection experiments (Drake & Pack, 2008; Suh *et al*, 2003). We hypothesize that an increase in intranuclear or perinuclear concentration of the Cas9 RNP could be obtained by combining PEI delivery with importin-mediated nuclear trafficking mediated by the nuclear localization signal (NLS) attached to the Cas9 protein (Mali *et al*, 2013; Cong *et al*, 2013)

In summary, HDR ranges from fractions of 1% ( Van Trung Chu *et al*, 2015; Lin *et al*, 2014; Miyaoka *et al*, 2016; Gutschner *et al*, 2016; Davis & Maizels, 2016) to ~60% (Richardson *et al*, 2016; Lee *et al*, 2017b), influenced by numerous factors including cell cycle (Gutschner *et al*, 2016), small molecule treatment ( Van Trung Chu *et al*, 2015; Lin *et al*, 2014; Li *et al*, 2017), upregulation of DNA repair proteins (Shao *et al*, 2017), siRNA/shRNA knockdown of NHEJ factors (Davis & Maizels, 2016; Van Trung Chu *et al*, 2015), dDNA design (Zhang *et al*, 2017; Richardson *et al*, 2016), timed protein degradation (Gutschner *et al*, 2016), locus, cell type and transfection method. However, a consistent and common approach has not yet been defined.

In this paper, we describe a proof-of-concept HDR system that may provide a simple and broadly applicable gene editing solution. The keys to our approach are combining 1) Cas9 engineered to co-localize with dDNA (Cas9 monoavidin:biotin dDNA) such that the dDNA is bound to the RNP at the time of DSB induction, and 2) application of dual nuclear localization modes (PEI and NLS) to ensure efficient delivery of the complex.

## Results

### Protein Engineering of Cas9 monoavidin (Cas9MAV)

We speculate that the current low HDR frequency by CRISPR/Cas9 is primarily due to the inefficient delivery of dDNA to a DSB site in the nucleus. Previous studies reported that colocalizing the dDNA to the DSB site using a chimeric crRNA:dDNA construct improved HDR (10%). While not requiring drugs, the improvement is not significantly better than cell cycle arrest (35%) (Lee *et al*, 2017b; Jinek *et al*, 2012). The crRNA:dDNA chimera used was exceptionally short in length (crRNA length 42, dDNA length 87), which could hinder dDNA annealing to the released proximal genomic DNA. In order to optimize delivery of dDNA to the DSB site, we conjugated dDNA to Cas9 through the high affinity avidin-biotin binding system. This system is facile to implement compared to multi-step chemical conjugation of sgRNA and dDNA chimeras, as biotin can be introduced to dDNA by a PCR primer extension or by purchase of a biotinylated donor DNA from a commercial vender. We hypothesized that HDR can be further improved by increasing the linker distance between the conjugated dDNA and the protein, thereby providing additional flexibility to capture the released genomic DNA. We therefore engineered a Cas9-monoavidin (Cas9MAV) construct, where Cas9 already containing dual NLS sequences (Lin *et al*, 2014), was further modified by addition of a 17-amino acid linker to a monoavidin domain at its C-terminus (Fig. 1A, Supplementary Figure S1). The rationale for choosing monoavidin was to maintain a 1:1 stoichiometry of the DSB site and dDNA being delivered, as opposed to streptavidin, which exists as a tetramer and could deliver up to four dDNA components.

**Figure 1.**
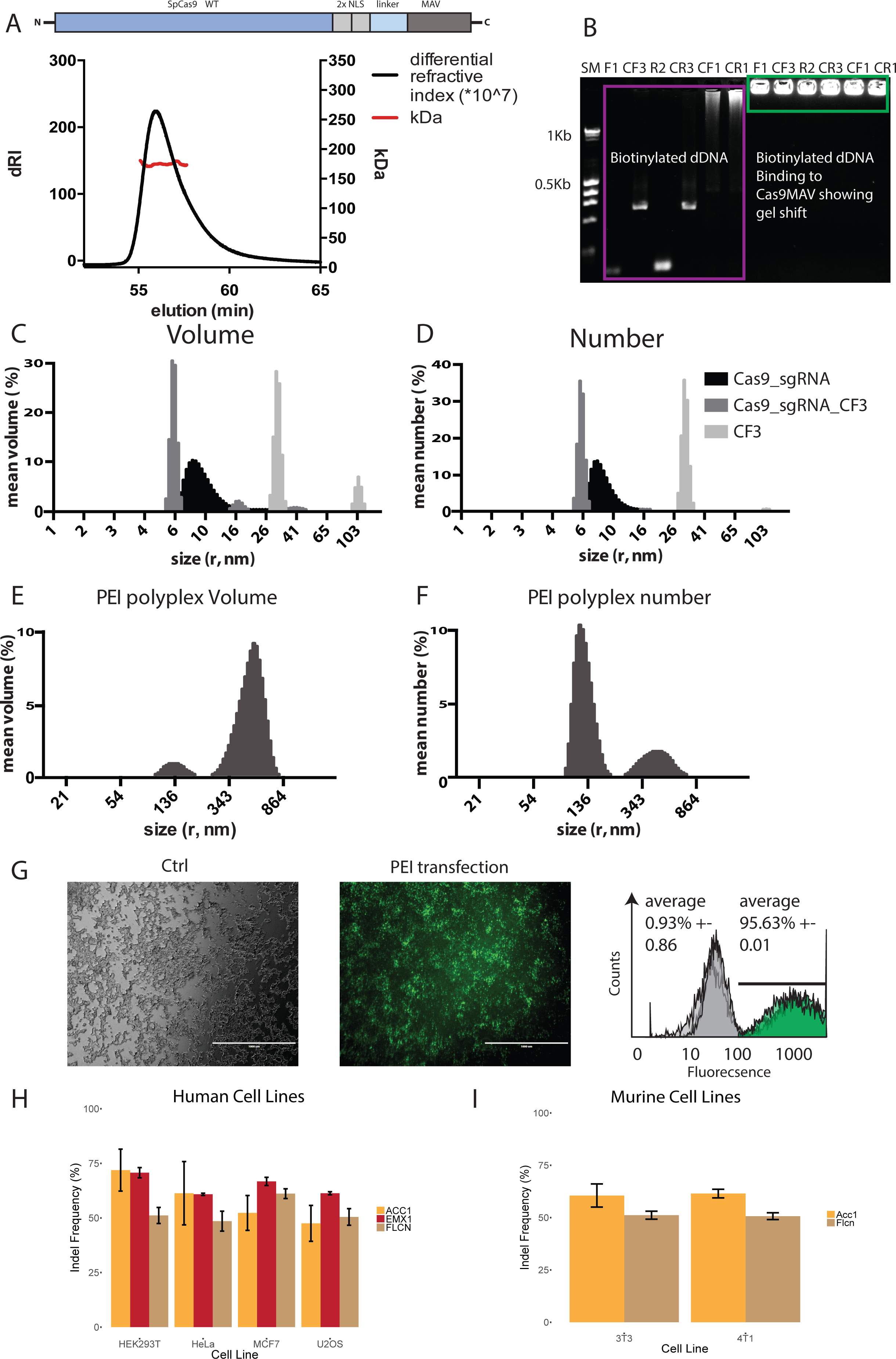
Characterization of the Cas9MAV protein **A**. SEC-MALS experiments were performed to determine the molecular weight and size of the modified Cas9MAV. The left y-axis shows the differential refractive index of the eluting protein peak (black), indicating perfect monodispersity. The calculated MW is 175.5 kDa and the measured MW across the peak, 174.5, is denoted on the right y-axis (red). An illustration of the Cas9MAV protein construct is included above the SEC-MALS figure. **B**. Migration of biotinylated DNA donors was tested on a 2% agarose gel in the absence (left, purple), or presence (right, green) of the Cas9MAV RNP. Abbreviations: SM (Size Marker), CF3 (CF3b), R2 (Ras2b), CR3 (CR3b), CF1 (CF1b), CR1 (CR1b). For donor structure and sequence, see Figure 2 and Table S2. **C**. Mean volume distribution of three consecutive samples of the complex formation assessed by DLS. The scattering of 2µM CF3 donor DNA (light gray) was firstly measured, followed by 2µM preformed Cas9MAV:sgRNA complex (black). Cas9MAV:sgRNA:CF3 RNP was measured (dark grey). **D.** Mean number distribution of the experiment shown in C. **E**. Mean volume distribution of the full polyplex. PEI was added directly to the preformed Cas9MAV:sgRNA:CF3 sample analysed in C and D and incubated for polyplex formation before being transferred back to the cuvette. DLS was measured as before, the peaks observed in C and D were undetectable, but two new populations of larger size are observed. The x-axis is adjusted accordingly. **F**. Mean number distribution of the experiment shown in E. DLS experiment was performed once with a minimum of 15 measurements per sample. **G**. Transfection efficiency of PEI. FAM-labelled PEI was transfected into HEK293T cells and fluorescence of control (left) and transfected (middle) cells was assessed using flow cytometry (right, biological replicates n=3, each repeat involves 5000 individual cell measurements). **H**. PEI:Cas9MAV creates double strand breaks. Human and murine cells were treated with PEI:Cas9MAV RNP without donor. Three genes EMX1, FLCN and ACC1 were targeted in human cells, while Acc1 and Flcn were targeted in murine cells. Each experiment for indel formation was evaluated as 3 biological repeats for each gene in each cell line.

Fig. 1A shows that Cas9MAV is monomeric in solution with a molecular weight (MW) of 174.5 kDa as determined by size-exclusion chromatography multi-angle light scattering (SEC-MALS) (calculated MW: 175.474 kDa). The hydrodynamic radius was measured to be 9.5 nm which is in accordance with the presence of the flexible NLS-linker-MAV tether in Cas9MAV and previously published Cas9 hydrodynamic radii (Carlson-Stevermer *et al*, 2017; Jinek *et al*, 2014; Nishimasu *et al*, 2014).

Fig. 1B illustrates an agarose gel shift assay with a subset of the biotinylated donors used in this study (Fig. 2A). In the absence of Cas9MAV (purple box) the dDNAs migrate according to their individual sizes, while in the presence of Cas9MAV (green box) no migration was detected due to the increase in molecular weight of the complex.

**Figure 2.**
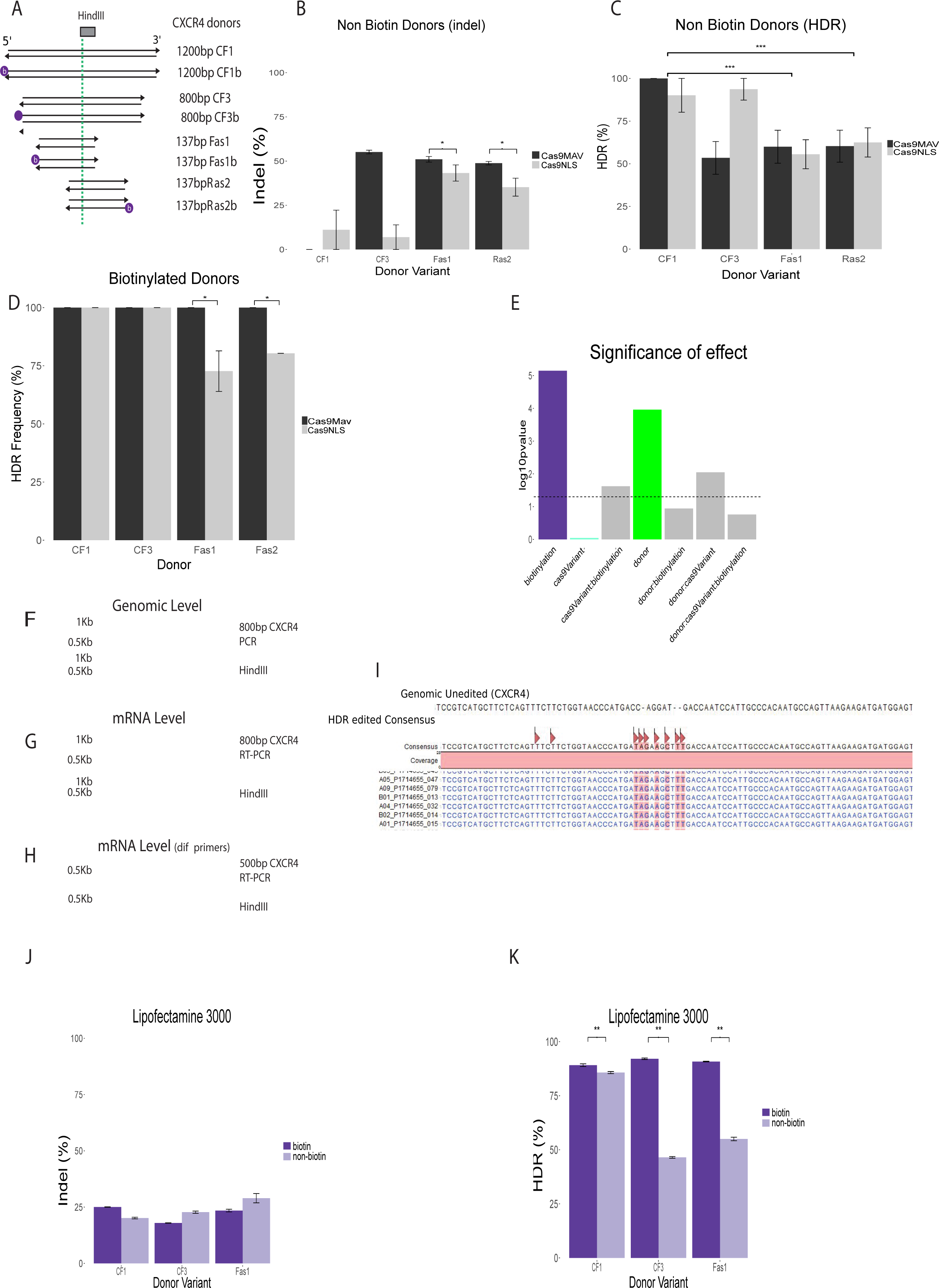
Summary of CXCR4 HDR Experimentation in HEK293T cells. **A.** Overview of the donor lengths and designs used, biotinylated and non biotinylated. “b” enclosed in a circle denotes position of biotin post PCR synthesis. Orientation of sense strand arrow indicated by 5’ and 3’ label. Green line denotes the central position and provides guidance as to the location of Stop codon (TAG) and HindIII sequence that follows. Lengths quoted in base pairs represents the total homology arm lengths used in each donor. The length of arrows either side of the dashed line indicates either equal homology arm length or asymmetric donor construction. **B** and **C.** Indel frequency (**B**) and its corresponding HDR experiments (**C**) with non-biotinylated donors. Significance was determined between the performance of Fas1/Ras2 donors and longer CF1 donors. **D.** Demonstration of HDR frequencies with biotinylated donors using both Cas9MAV and Cas9NLS. **E** summarizes ANOVA results, considering donor design, biotinylation and usage of Cas9NLS and Cas9MAV. The two significant factors are donor design and biotinylation of dDNA with Cas9MAV, which outperform Cas9NLS. **F.** 800 bp amplicon expressed from genomic DNA and the commensurate HDR derived HindIII cleavage. **G** and **H.** RT PCR products (800 bp and 500 bp) and their HindIII cleavage. At both genomic and mRNA levels the edit was shown to persist. Each experiment was repeated with biological replicates of 12 initial samples. 1 negative control, 11 successful RNA/cDNA reverse transcription gave 8 successful RT-PCRs. Fig. **2G** shows the first 3 lanes were negative amplification to be specific (lanes numbered 4, 5, 6, 7, 8, 9). Fig. **2H** shows the second RT-PCR gel where 8 amplifications were successful and resulted in 8 digests (lanes numbered 4, 5, 6, 7, 8, 9). **I.** Summary of Sanger sequences from different Cas9MAV experiments with the 4 biotinylated donors illustrated in A. Base sequence modification caused by insertion of donor DNA is marked with red triangles. **J** and **K** detail the indel and HDR frequencies using lipofectamine 3000 and biotinylated/non-biotinylated CF1, CF3, Fas1 donors (see Figure 2A). **J** details the indel average percentage evaluated by T7 endonuclease assay for non-biotinylated/biotinylated donors. **K** represents the results of HindIII digest HDR assay. All plots in this figure represent a minimum of 3 biological replicates. In all bar plots, error bars represent standard deviation. Statistical analysis was performed using one way Anova and Tukey test with p-values: * = 0.05, **=0.01, ***=0.001.

### Characterisation of PEI:Cas9MAV Polyplexes

Before attempting colocalization intracellularly, formation of a Cas9MAV complex bound to biotinylated dDNA was first demonstrated *in vitro*. Analysis by dynamic light scattering (DLS) (Fig. 1C and 1D) established that near complete conjugation of dDNA to Cas9MAV RNP was achieved.

DLS measures light scattering generated by particles representing size, distribution and polydispersity. Sample mean intensity distribution can be transformed to a mean volume distribution (sample population spatial occupancy), and a mean number distribution (number of particles per population per volume). Fig. 1C and 1D shows that Cas9MAV:sgRNA is monodisperse with a 9 nm radius (black histogram) correlating well with the SEC-MALS results of Cas9MAV alone. The dDNA tested was the 800 bp long CF3b (5’ biotinylated) and it was determined to have an approximate radius of 30 nm (Fig. 1C and 1D, light grey histogram). Next, preformed Cas9MAV:sgRNA:CF3b complex (1:1:1 molar ratio) was measured, showing that CF3b was able to complex completely with Cas9MAV:sgRNA (Fig 1C and 1D, dark grey histogram), as no residual free CF3b dDNA could be detected upon RNP addition. It was observed that the radius of the final Cas9MAV:sgRNA:CF3b complex was smaller, with a hydrodynamic radius reduced to 6 nm, although a small population was observed at ~ 16 nm. In conclusion, sequential RNP assembly is confirmed by the absence of the free macromolecule populations (i.e. the CF3b and Cas9MAV:sgRNA populations disappear when sgRNA:Cas9MAV:CF3b forms).

DLS was also used to validate polyplex formation by studying encapsulation of the RNP complex by PEI (Fig. 1E and 1F). PEI was added directly to the Cas9MAV:sgRNA:CF3b RNP sample described above and incubated at room temperature for 10 min before DLS measurements. The Cas9MAV:sgRNA:CF3b population was no longer detectable, indicating that it formed a polyplex with PEI. By intensity and volumetric measurements, we observed two species of polyplex at 134 nm and 530 nm radii. Considering the number of species in each population, PEI polyplexes of 134 nm radius were the most prevalent. Thus, based on this radius and prior knowledge of PEI, it is most likely that calnexin-mediated and macropinocytosis mechanisms (Khalil *et al*, 2006) are employed by the Cas9MAV:sgRNA:dDNA:PEI polyplex to cross the cell membrane in transfection experiments, since these mechanisms are more compatible with objects of 130 nm.

### High transfection efficiency with PEI:Cas9MAV

Successful HDR is comprised of several steps - polyplex formation, cellular uptake, transport to the nucleus, creation of DSB, and correct repair with dDNA. High transfection efficiency must be obtained for the proposed Cas9MAV system to function. To track transfection of the PEI:Cas9MAV polyplex, we used fluorescent PEI. Fig. 1G shows bright field and fluorescent light microscopy of transfected cells, respectively, with FAM-labelled PEI:Cas9MAV without dDNA (see Supplementary Methods). This confirms that transfection has occurred qualitatively, since the cells are positive in the green fluorescence imaging channel. Quantitative measurements using flow cytometry showed that 95% of cells were transfected by the FAM-labelled PEI:Cas9MAV polyplex (Fig. 1G, right). Having demonstrated successful cellular uptake, we next investigated nuclease activity in multiple cell lines.

Six cell lines (Human: HEK293T, MCF7, HeLa, U2OS and Mouse: 4T1, 3T3) were chosen to reflect both easy-(HEK293T, HeLa) and difficult-to-transfect (4T1, MCF7) cells, two species, and a variation in tissue of origin/disease (fetal kidney, breast cancer, cervical cancer, osteosarcoma, fibroblasts). To examine multiple loci, RNPs were formed complexing Cas9MAV with sgRNA targeting FLCN, ACC1 and EMX1 for human cell lines (Fig. 1H) and sgRNAs targeting Flcn & Acc1 for murine cell lines (Fig. 1I). NHEJ indel frequency was determined by a T7 endonuclease assay after transfection with PEI:Cas9MAV without dDNA in all cell lines. Indel frequencies of 50-70% were found across the human cell lines (Fig. 1H). In murine 4T1 and 3T3 cell lines, indel frequencies of 50-60% were observed in the two loci tested (Fig. 1I). These results demonstrate that the PEI:Cas9MAV could be successfully transfected and induce indels in other cell lines besides HEK293T.

### High HDR frequency in combination of PEI and Cas9MAV-dDNA complex

To determine whether Cas9MAV can mediate efficient HDR editing and to investigate the impact of donor design on Cas9MAV editing efficiency, we outlined a simple comparison experiment at the CXCR4 locus, focusing on four double stranded biotinylated and non-biotinylated dDNA designs (Fig. 2A, Supplementary Table S2). Each dDNA was designed to introduce a HindIII restriction site within 5 bp of the DSB site. The intention was to explore the design rules of the length of HAs for double stranded dDNAs. We generated HA lengths of 600bp (CF1), 400bp (CF3) and short asymmetric donors (Fas1 and Fas2). The latter are possibly sub-optimal designs as their HAs are significantly truncated. Indels were detected by T7E1 DNA mismatch detection assay and HDR was analyzed by HindIII restriction fragment length polymorphism assay (RFLP) (Fig 2B and 2C). For Fas1 and Fas2 dDNA, indel percentages of 40-60% persisted (Fig. 2B) and conversely, HDR percentages of 50-60% were observed (Fig. 2C), for both Cas9MAV and Cas9NLS. CF1 and CF3 dDNA averaged only 5% and 24% indel formation with Cas9NLS, respectively, and 0% and 60% with Cas9MAV. HDR percentages for CF1 and CF3 with both Cas9NLS (85-90%) and Cas9MAV (100% and 50%, respectively) were roughly the inverse of indel percentages. Thus, shorter non-biotinylated dDNAs generally result in lower HDR percentages, suggesting that they are less likely to form stable, hybridised species with genomic DNA than longer dDNAs.

Improved HDR percentage was observed upon biotinylation of the donor constructs, including the shorter Fas1b and Ras2b (biotinylated version of Fas2) (Fig. 2D). Near 100% HDR was found across the four dDNAs tested with Cas9MAV, illustrating that the observed HDR improvements are a consequence of the combined action of the Cas9MAV protein and transfection via PEI. Cas9MAV results in smaller improvements in the long donors, since they are already close to optimal. However, we can see substantial improvements in the short donors. Fig. 2E summarizes the overall significance of each factor, as calculated by ANOVA. It shows that donor design plays a significant role (Green) as does biotinylation (Purple), which we interpret as colocalization by Cas9MAV over all donors. At the CXCR4 loci, utilization of biotinylated donors and Cas9MAV contributes a 20% improvement in HDR rates in comparison with Cas9NLS. Biotinylation when using Cas9NLS does not improve the HDR rates (Tukey HSD tests, see Supplementary SD5)

Next, we wished to validate the performance of the biotinylated dDNA and Cas9MAV by investigating whether the genomic edits persisted at the mRNA level. Using Fas1b to edit cells, inserts were validated using its associated HindIII site (Fig. 2F-H). Genomic DNA was degraded by DNAse1 before reverse transcription (see Supplementary Methods). Two primer sets for two separate extractions demonstrated the presence of the edit on the mRNA level (Fig. 2G and 2H). This enabled us to determine by cDNA derived PCR products, that restrictive cleavage was preserved and transcribed to mRNA, demonstrating the persistence of the edit. Furthermore, the edits persist after significant dilution and re-growth of cells post passaging (Supplementary Figure S2). Samples edited with CF1, CF3 and biotinylated CF1 and CF3 were prepared for Sanger sequencing and all samples returned the correct HDR insert sequence (Fig. 2I).

### PEI Transfection and Cas9MAV impacts HDR: A comparison with Lipofectamine 3000

Central to our system are PEI transfection and the monoavidin modification, offering a nuclear trafficking approach and dDNA co-localization at the DSB site, respectively. Nucleofection was shown to result in comparable indel formation with Cas9MAV and Cas9 NLSat CXCR4 and EMX1 loci (Supplementary Fig. S3). This result suggests that nucleofection renders the NLS sequence redundant. Early experiments using the lipofection technique produces cytosolic delivery for DNA payloads (Akita *et al*, 2004). Consequently, we analyzed transfection by Lipofectamine 3000 as a substitute method for PEI, but without the nuclear location potential and lacking microtubule transport to the perinuclear space (Cardarelli *et al*, 2016). Lipofection shares many characteristics with PEI, such as payload (DNA/protein), protection from metabolic degradation, high transfection rates and low cytotoxicity (Supplementary Fig. S4). Lipofectamine and its derivatives such as CRISPRmax have shown high transfection and HDR editing (~20%) (Yu *et al*, 2016). However, lipofection lacks nuclear localization and microtubule transport, and thus provides an ideal comparison transfection method to PEI.

Fig. 2J and 2K show the results of transfection experiments using Lipofectamine 3000 and Cas9MAV for editing the CXCR4 loci with the same donors as above (Fig. 2A). Fig. 2J compares indel percentages between biotinylated and non-biotinylated donors. It is observed that indels decline where biotinylated donors are used, with CF1 being an aberration in this instance with persistent indels. As described above, decreased indel formation suggests increased HDR. Fig. 2K confirms that biotinylated donors outperform non-biotinylated donors (CF1:CF1b = 85% to 89%; Fas1:Fas1b = 55% to 90%; CF3:Cf3b = 46% to 92%). Figure 2K also shows again that the donor design plays a substantial role, with CF1 as the optimal donor, achieving very high percent without biotin. Fas1:Fas1b show a greater degree of improvement. Thus, the difference between the biotinylated and non-biotinylated donors confirms that dDNA colocalization with Cas9MAV increases HDR. Importantly, transfection with Lipofectamine 3000 does not achieve the same HDR percentage as is seen in cells transfected with PEI (Fig. 2D), suggesting but not validating that the nuclear localization potential of PEI plays a significant role. Further mechanistic evaluation is beyond the scope of this paper, but both PEI and monoavidin modifications have statistically relevant effects on the HDR achieved in this work.

### Traffic Light Reporter (TLR) Assay

Given the high HDR percentages observed in the CXCR4 locus, we wanted to demonstrate the versatility of our PEI:Cas9MAV system by applying it to other loci. CXCR4 locus is usually associated with high indel percentage without a donor, and thus has the potential for high HDR percentage with dDNA. It was important not to choose loci and commensurate sgRNAs with performance equivalent to CXCR4, as this would not give a clear evaluation of general application of the system. We chose EMX1 and a Traffic Light Reporter (TLR) system (Certo *et al*, 2011), to obtain a more wider appraisal of performance. The TLR system is a cassette inserted into a safe harbour locus that expresses red fluorescent protein, when HDR occurs the template introduces a frameshift that converts the protein to green fluorescent protein.

The TLR assay appeared an appropriate choice where DSB are low, since indel percentages observed by the TLR assay are usually very low (12%)(Robert *et al*, 2015). Evaluation of the assay was performed using the Cas9NLS plasmid as an indel-forming positive control (Supplementary Fig. S5). At the 96 hr time point, complete repair of DSBs will have occurred, and there has been sufficient time for the Cas9 plasmid control to reach a measurable signal (RFP expression) for flow cytometry. NHEJ frequency of 17.1% was observed at 96 hrs for the Cas9 plasmid control, similar to previous NHEJ percentages with this assay (Robert *et al*, 2015). When using biotinylated donors and PEI:Cas9MAV, we observed a decline in NHEJ (from 12% to 0%) and HDR of 1-3% with 5’b and 3’b dDNA variants compared to control (Fig. S5) at 48 hr post transfection (Supplementary Table S3). We observed that it was possible to detect the mixed NHEJ/HDR population in cells edited with the PEI:Cas9MAV system with both flow cytometry (Supplementary Figure S5) and fluorescence microscopy (data not shown). At 96hr, NHEJ increases to 0.23-0.61% and HDR percentage for 5’b, 3’b and dual biotinylated 5’3’ donors with the PEI:Cas9MAV was 20-32%. In conclusion, NHEJ was a minor contributor to edits with less than 2% for all biotinylated donors. Representative examples of Cas9 plasmid control without donor (96 hrs) and PEI:Cas9MAV with 5’b donor (96 hrs) are given in Fig. 3A and 3B, respectively. Full flow cytometry plots and results are presented in Supplementary Figure S5.

**Figure 3.**
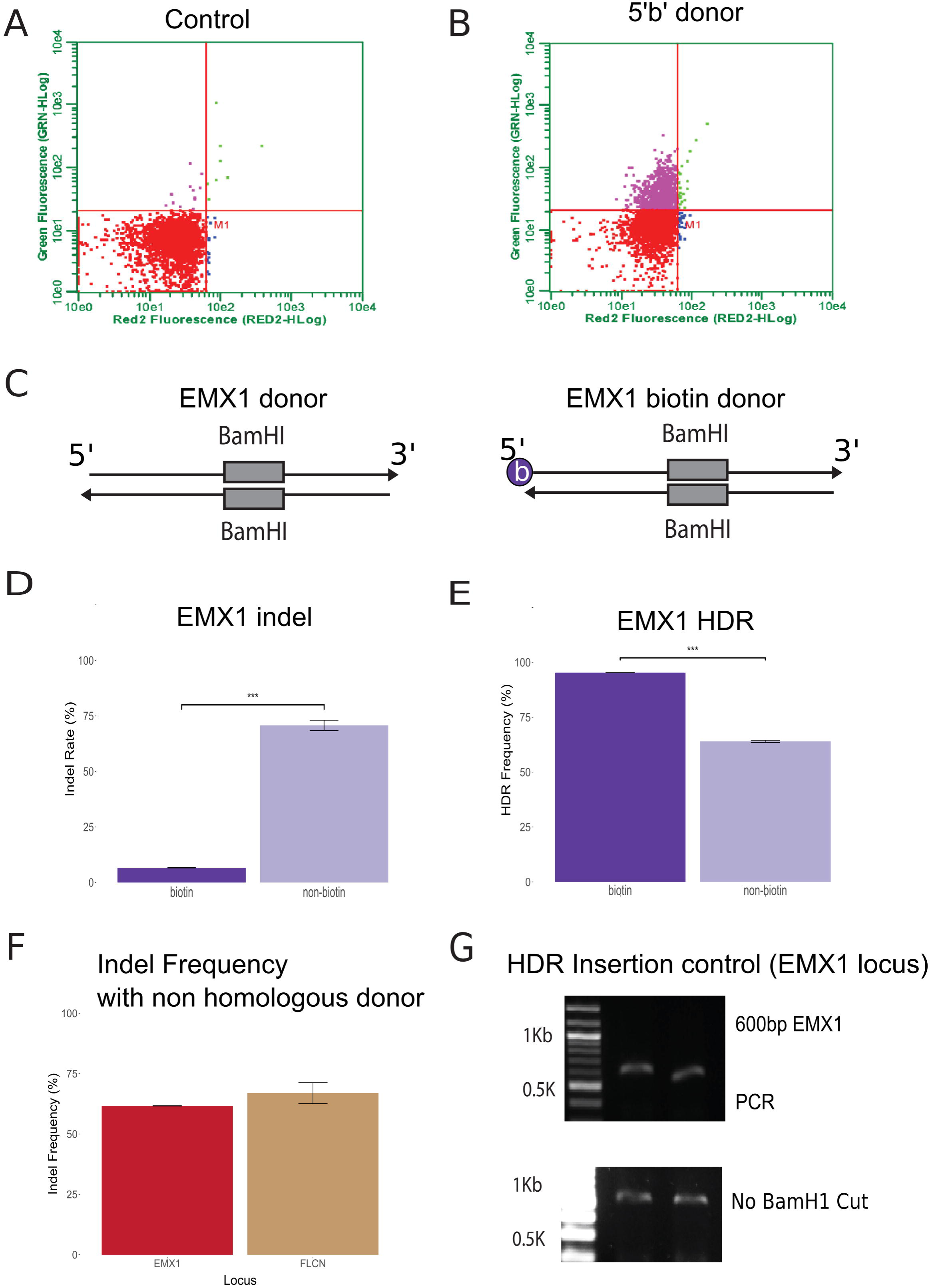
HDR percentages using Traffic Light Reporter and the EMX1 locus. **A** shows the control population used to gate for flow cytometry, and **B** shows HDR caused by using Cas9MAV with donor 5’b, as an indicative example of HDR observed as occurring at the 96hr time point, occurring in the TLR assay with PEI:Cas9MAV. The X axis represents red fluorescence (NHEJ), while the Y axis represents green fluorescence (HDR). Further analysis of TLR assay at 48hrs and 96hrs is provided in Supplementary Figure S5. **C** shows the two donor designs for EMX1, where the inserted BamH1 site is in the center of the designed sonor, between equal homology arms of 200bp at the EMX1 locus. Donors were generated with matched non-biotinylated and biotinylated donors and PEI:Cas9MAV. Total donor length was 400bp (+ BamH1 & quadruple stop codon repeat) and contained a BamH1 site for validation of insertion. Biotinylation at the 5’ end of the forward primer is denoted by ‘b’. **D** and **E** show indel and HDR frequencies with biotinylated and non-biotinylated donors. T7 (**D**) and HDR (**E**) assay samples are summations of a minimum of 3 biological replicates. **F** considers indel frequency at EMX1 and FLCN loci after transfection with a mismatched biotin donor for CXCR4. Indel frequency remained constant. **G** is an initial investigation as to whether a donor not matched to genomic sequence can insert at a locus cleaved by Cas9MAV using our method. EMX1 loci PCR amplicon which does not form digest, showing mismatched donor was not inserted. All plots in this figure represent a minimum of 3 biological replicates. In all bar plots, error bars represent standard deviation. Statistical analysis was performed using one way Anova and Tukey test with p-values: * = 0.05, **=0.01, ***

### EMX1

The HDR percentages achieved by our drug-free system in both CXCR4 and TLR experiments represented a substantial improvement on previous pharmacological interventions ( Van Trung Chu *et al*, 2015; Certo *et al*, 2011). In keeping with the trends observed for the CXCR4 gene locus (Fig. 2), we surmise that EMX1 indel frequencies are a metric for sgRNA quality and genomic locus accessibility. Based on observed indel of 65-70% (Fig. 1H), we predicted lower HDR than those observed for CXCR4, but greater than TLR using the PEI:Cas9MAV system. We used a lower quality donor compared to the CXCR4 locus, comprised out of 200bp equal HAs on either side of a 6-repeat stop codon sequence and BamHI site. The experiment assessed two donors, biotinylated and non-biotinylated (Fig. 3C).

The non-biotinylated EMX1 donor resulted in HDR percentage of 60% with high (~70%) indel formation; these results sum to more than 100%, reflecting the semi-quantitative inaccuracies of gel-based assessments and suggesting mono-allelic editing, partial incomplete HDR (leading heteroduplex and indel assignment by T7endonuclease assay) and reflecting the semi-quantitative inaccuracies of gel-based assessments. This observation correlates with the lower HDR frequencies generated by short double stranded donors, as used in our CXCR4 investigations (Fas1/Fas2 137 bp) and in the TLR study (111 bp total). Use of biotinylated EMX1 donor results in indels dropping to less than 10% and HDR increasing to 90% (Fig. 3D and 3E), further confirming that HDR improvements can obtained with Cas9MAV protein for colocalization of dDNA.

We also wished to consider the impact of donor DNA being inserted randomly where the Cas9 DSBs were generated. A simple experiment was performed where the sgRNA for EMX1 was mismatched with a biotinylated CF3 CXCR4 donor. EMX1 retained its capacity for indel generation (Fig. 3F) and the absence of insert was validated as HindIII digestion did not result in cleavage (Fig. 3G). While not a complete proof of non-insertion where Cas9MAV cleaves DNA, it offers some indication that non-matched donors do not result in HDR.

### NGS sequencing Confirmation of Editing Frequencies

To consider HDR at a greater depth of sequencing we designed an Ion Torrent sequencing approach. We took 7 CXCR4 sample experiments covering HDR (with and without biotinylated donor), and unedited controls (Supplementary Tables S4, S5). Fig. 4A and B present an overview of the libraries sequenced, which are detailed in Supplementary Table S4. Untransfected HEK293T cells (FControl) returned a clear unedited population of sequences, replicating the canonical genomic sequence for CXCR4. The untransfected controls cells showed 1.3% of sequences matching the HindIII site, giving us an approximation of sequencing errors in this locus, this level of background was also observed in other HDR systems. Our principal interest was to evaluate the sequences returned for complete edits (full insertions of donor sequence) and partial edits. HDR experiments (FBest, F60, BC4, BC5 and BC6), show that HDR has occurred at a 90% or greater (Fig. 4A and 4B). The CXCR4 library with Cf1b donor (FBest) is a good indication of correlation with gel-based analysis given in Fig. 2. These results suggest that suppression of indels and high HDR percentage was achieved for the complete insert, with less than 5% being partial inserts/indels. In this case, the restriction digest provides a good guide for donor design performance. When Fas1, a non-biotinylated asymmetric donor was analysed (F60 library), we observe an unedited fraction persisting, but at a lower percentage (~2%) than was estimated from the restriction digest (Fig. 2C).

**Figure 4.**
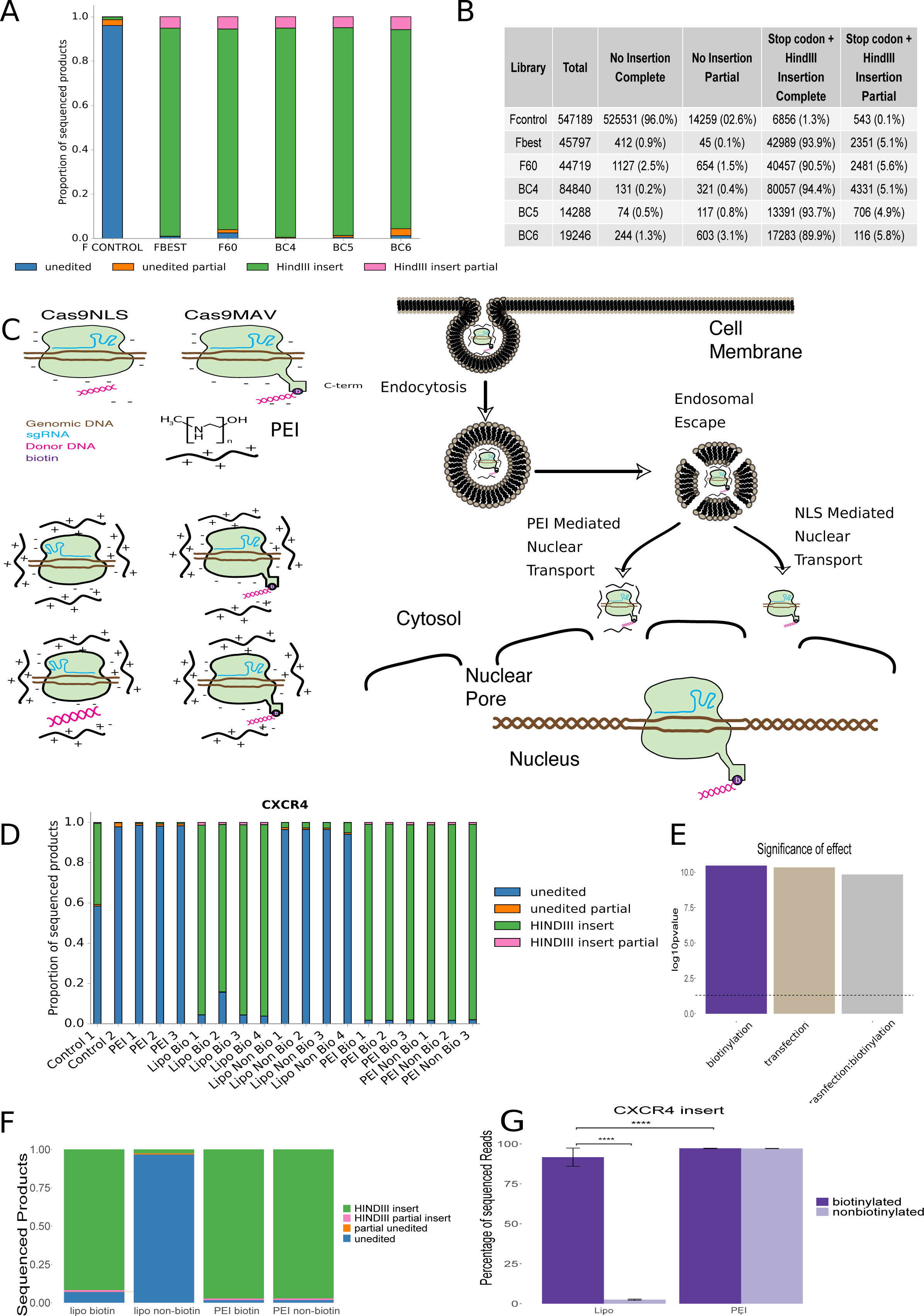
Deeper consideration of the PEI:Cas9MAV system towards a proposed model. **A** shows a high level evaluation of library from the first Ion Torrent Sequencing. Each sequence was subdivided firstly on the basis of unedited and edited sequences and grouped by barcode to give total reads per library. Library name and commensurate donor variant used with PEI:Cas9MAV are detailed in Supplementary Table S4. NGS provides a more quantitative method of evaluating HDR than either TLR or digest assays, as direct sequence analysis can provide information relating to the knock-in variation or partial insertions. **B** is a table describing the complete and partial insertion sequence numbers (with percentages) from Ion torrent sequencing analysis. Total reads implies total reads per library. Complete refers to exact alignment to either no insert control sequence or inserted HDR sequence. Partial refers to alignment to either control sequence or inserted HDR sequences with **C** Schematics of the hypothetical PEI:Cas9MAV system based on the experiments conducted. Two variants of canonical Cas9 are considered. Cas9NLS is a commonly used variant enabling nuclear localization, Cas9-NLS-monoavidin (Cas9MAV) is a new variant developed for this work containing a C-terminal peptide linker and monoavidin domain. The cationic polymer used to form polyplexes is represented in its general monomeric form and in cartoon lines with (+) representing charge. Using both variants, Cas9NLS can exist in association in a PEI polyplex with or without DNA as there is no physical link. Cas9MAV applies biotin linkage to bind donor DNA with the MAV domain, and therefore Cas9MAV should always exist with the donor. It is hypothesized that PEI:RNP could enter the cell via endocytosis and undergo endosomal escape. Nuclear transport could happen via two mechanisms: PEI mediated nuclear transport, or entrance by the C-terminal nuclear localization sequence recruiting importins for internalization. Once in the nucleus, Cas9 and donor are available to perform HDR. **D** is a high level evaluation of library from the second Ion Torrent Sequencing, which included sequencing the lipofectamine results. Each sequence was subdivided firstly on the basis of unedited and edited sequences and grouped by barcode to give total reads per library. Library names and commensurate donor variants used with PEI:Cas9MAV are detailed in Supplementary Table S6. **E** summarizes of ANOVA results, considering donor biotinylation and transfection method (PEI vs Lipofectamine 3000). Both factors are significant, and are significant in combination (all p-values smaller than 1e-10). **F** summarizes the sequencing results for PEI and Lipofectamine 3000, using both biotinylated and non-biotinylated donors. The order is Lipofectamine non-biotinylated, Lipofectamine non-biotinylated, PEI non-biotinylated and PEI biotinylated. **G** summarizes the percentage of HindIII insert under each condition. All plots in this figure represent a minimum of 3 biological replicates. In all bar plots, error bars represent standard deviation. Statistical analysis was performed using one way Anova and Tukey test with p-values: * = 0.05, **=0.01, ***, **** = <1e-10

Initial NGS sequencing was extended to investigate other experimental conditions: CXCR4 using lipofectamine with biotinylated and non-biotinylated donors in comparison to PEI transfection. Common to all experiments sequenced was the use of the shortest donor construct for the CXCR4 loci (137bp). The purpose of selecting these experiments for sequencing was to look at the concept proposed in Fig. 4C, that PEI transfection provides an alternative means for nuclear location of RNP:dDNA (biotinylated and non-biotinylated). It was hypothesised that lipofection, which does not locate to the nucleus, would therefore abrogate or reduce HDR frequencies, as previously seen in the restriction digests in Fig. 2K.

Fig. 4D shows the sequencing results for each of the libraries covering these conditions. Full library descriptions and table are present in Supplementary Table S6. Confirmation of the restriction digests and the first sequencing results was observed for PEI (biotinylated and non-biotinylated dDNA) experiments, with HDR read percentage at ~94%. A summary figure is presented in Figure 4F.

ANOVA (Fig 4E) and Tukey analysis (see SD5 supplemental), showed that transfection method is a definite factor in determining HDR. Using lipofection with non-biotinylated dDNA, HDR percentage declined to 2.5-5% of total reads, while using PEI and non-biotinylated donors maintained an HDR percentage in excess of 90% (Fig 4G, p-value < 1e-10). This comparison indicates PEI potentially has a role in nuclear location of donor, worthy of further investigation (Fig. 4F). Lipofection with non-biotinylated donors is validated as a significant abrogation of HDR. Biotinylated donors resulted in HDR percentage in excess of 90% for both lipofectamine and PEI, indicating the importance of Cas9MAV:biotin dDNA conjugation as well as the nuclear localization by NLS. Note that biotinylation in combination with lipofectamine showed a greater increase than biotinylated dDNA with PEI, simply because the base success rate with PEI is already high. Further studies on donor nuclear localisation and biotinylated donor forms will be required to validate our hypothesis about mechanism of action (Fig. 4C), which is outside the scope of the current work. In particular performance at other loci, cell lines and with larger donor insertions will be key to forming a nuanced understanding of RNP:donor constructs.

## Discussion

We demonstrated the potential for a cationic polymer delivery of RNP as part of a streamlined and efficient Cas9 HDR method. Biotin-avidin conjugation of the Cas9 RNP with a donor DNA and PEI dependant delivery to cells appear to be contributing factors in the improvement of HDR percentage observed at the CXCR4, TLR and EMX1 loci. Our HDR rates matched and surpass the current best practices, without using drugs, and resulting in a streamlined and simple method.

The ANOVA results show that there are several potential factors that influence HDR success rate. The first factor is donor design and second factor is transfection method, where PEI results in quite high HDR, even with potentially sub-optimal donors. The third factor is biotinylation of the donor together with our Cas9MAV system, and this is very effective when using sub-optimal donors and/or when using transfection methods such as lipofectamine. It does contribute to improvement when using PEI and optimal donors, but since the results are much closer to 100% to begin with, our method cannot improve HDR beyond 100%. As such, this suggests the possibility of using short donors, conforming to Richardson rules (Richardson et al), which are quite easier and cheaper to synthesis as a GBLOCK, and resulting in HDR success rates that rival or surpass donors with longer homology arms, which are more complicated to synthesize.

Cationic polymers generated high NHEJ at other loci (FLCN, ACC1) when dDNA is not present. Further work is required to unravel the contributions of biotinylated donors and nuclear localization in cell lines, and how principles translate to stem cells and *in vivo* settings. Additional studies should be performed on RNP delivery relating to polymer derivatization and polyplex size (Ogris *et al*, 1998). These polyplex properties have been shown as key determinants of DNA transfection in different cell types and in-vivo.

The monoavidin:biotin conjugate appears to relax further the design rules that usually operate for template donor during HDR (Fig. 2). In previous studies, the necessity for double stranded dDNA with HAs of 400-600 bp has been required to achieve high HDR frequencies (Lin *et al*, 2014; Zhang *et al*, 2017). Using Cas9MAV, biotinylated asymmetric short double stranded dDNA (137 bp, HA 37 bp and 93 bp, respectively) can achieve equal efficiency of HDR, mitigating the suboptimal design. This may be useful for simpler synthesis of DNA donors.

PEI provides an alternative to nucleofection/electroporation techniques and is simple to apply, but for completeness Cas9MAV RNP was delivered by nucleofection (Supplementary Fig. S3). Nucleofection may lead to potential problems in translating the transfection method into *in vivo* studies, since animals have a long history of electroporation (Adachi *et al*, 2002), but human studies are still under development (Jiang *et al*, 2015) for viral and non-viral transfection methodologies.

To codify the experimental results, the combinations of loci:sgRNAs chosen were loosely termed the ‘good’, the ‘bad’ and the ‘ugly’, by indel frequency without HDR. The ‘good’ refers to a locus where the sgRNA scores highly in design programs and indel frequency is in excess of 60-80% as indicated by T7E1 endonuclease assay. We hypothesize that for HDR, a consistently high occurrence of DSBs is required with high recognition of target by sgRNA. We acknowledge that T7E1 assays cannot reflect conservative DSB repairs, but offer only an approximate measure of cut efficiency. The ‘good’ approximation refers specifically to the performance at CXCR4. The ‘bad’ is a locus (EMX1) where the indel frequency is around 60% denoting lower DSB frequency or higher conservative DSB repair. In case of the ‘bad’ locus, success was determined by achieving HDR above 60% to be consistent with the hypothesis that indel and HDR frequencies in this system invert. The ‘ugly’ is an HDR assay using the traffic light reporter system (Certo *et al*, 2011), that in previously published examples exhibits very low HDR percentage ranging from 1-4%. Correspondingly, TLR experiments show comparatively low NHEJ at 12% or less, even with Cas9 plasmid systems, reflective of the locus:sgRNA combined inefficiency. In this case, the determination of success achieving HDR in excess of 4% without drug treatment, siRNA knockdowns, or other interventions (Robert *et al*, 2015). In the three loci tested, we observed high percentages of HDR occurrence and suppression of NHEJ, with varying degrees of HDR between the loci due to experimental factors.

We caution that any Cas9 experiment’s final editing efficiency will be governed by a combination of multiple factors that include: loci accessibility (condensed/uncondensed chromatin), sgRNA quality (sequence recognition & DSB efficiency), donor sequence/size (geometry, size. It can not be assumed that co-localisation can produce high HDR occurrence at all loci, the TLR experiment is an exemplar, demonstrating that a poor locus:sgRNA combination, resulting in maximal HDR percentage of only 25%.

Our approach has drawbacks, where biotinylated donors contribute to overall experimental cost, and the impact of placing the donor DNA on a flexible mono-avidin linker cannot be fully validated other than by further experimentation. Furthermore, little mechanistic understanding that can be contributed in this work, regarding why HDR is favoured and the model proposed in Figure 4 is purely deductive reasoning. It is hoped that future DNA repair experiments will provide context.

The ability of the Cas9MAV system to remove many of the donor design constraints (length, asymmetric, single/double strand, HA length), offers an intriguing opportunity we hope to explore by using other loci, as well as varying donor DNA size and insert size. There are few general rules for donor constructs. Richardson et al (Richardson *et al*, 2016) provide guidance that is reproduced in other papers for short donors using ssODN, while larger constructions remain ill defined and the effect of biotinylation on HDR of larger inserts remains to be explored.

Our results do not stand in isolation. Recent publications exploring the role of donor localization to the nucleus and importantly, DSB site (Carlson-Stevermer *et al*, 2017; Ma *et al*, 2017), offer support that our system is affecting the mechanisms of DSB repair. It is uncertain at this point which repair pathway and protein effectors of repair, such as DNA-ligase, Ku heterodimer, Rad51/52, are involved and at what stage. Fig. 4C summarises our current model, focused on the benefit derived from biotin mediated donor localization with two nuclear delivery approaches.

In contrast to other CRISPR/Cas9 methodologies involving plasmid or mRNA delivery, our system does not require the cell to transcribe sgRNA and translate protein, which can impede RNP intracellular assembly. By coupling donor to Cas9, we remove the issues of a freely diffusing donor (plasmid or linear) and compartmentalisation of the donor in the cytosol. If the donor is not present, it is more likely that canonical DSB repair (NHEJ and conservative blunt end repair (Bétermier *et al*, 2014)) will occur rather than HDR. Recent work using aptamer:streptavidin localisation introduced ssODN to nuclei (Carlson-Stevermer *et al*, 2017), achieving ~6% HDR. This rate is lower than what could have been achieved with extracellular RNP assembly and hence an increased concentration of functional RNP:donor conjugates within the cell. In future work we hope to extend the transfection principle and Cas9 methodology to difficult-to-transfect cells other than MCF7 and 4T1.

## Materials and Methods

### Cell culture

HEK293T, MCF7, 3T3, HeLa and U2OS cell lines were all cultured in DMEM supplemented with 10% FBS, 100 U/mL penicillin, and 100 U/mL streptomycin and were maintained at 37°C and 5% CO2. The exception being 4T1 which was cultured in DMEM with 10% FBS, 10mM HEPES, 1mM Sodium Pyruvate, 36mM NaHCO_3_, 5 µg/mL amphotericin, 0.5% gentamicin and 1% L-glutamine. Media, trypsin and FBS were supplied by Wisent. Cells were kept at low passage for experimentation, not exceeding 10 passages before starting fresh cultures from frozen stocks.

For transfections, seeding density was 3×10^5^ cells per 6 well and 1×10^5^ for 12 well plates, prepared on the day prior to PEI transfection. Confluency of 60-70% at time of transfection was our objective, so for fast growing (HeLa) or larger cells (U2SO), seeding densities of 1×10^5^ and 0.8×10^5^ respectively were used for preparation for 6 well plates.

### Cell Viability

Cell viability was determined by trypan blue staining in a 1:1 v/v ratio to 5µl sample of harvested cells. Staining was verified by Countess II or visually by haemocytometer. All cells were maintained at 97% viability post transfection.

### sgRNA

Full sequences and synthesis by in vitro transcription are described in Supplementary Methods.

### PEI preparation

100mg of linear PEI (polyscience) was dissolved in 100 ml nuclease free milliQ water. Solution was magnetically stirred for 4hrs at room temperature in a sterile Duran bottle. Once the polymer was dissolved, pH was adjusted to pH 7.4 using concentrated HCl. Secondary amine protonation was assessed by ninhydrin assay by taking aliquots. Solution was filtered (0.22um) and aliquoted into 1.5 ml eppendorf tubes and stored at −20°C. Aliquots were removed as required for experiments and stored at 4°C for polyplex formation, with a maximum usage period of two weeks. PEI fluorescent labelling with carboxyfluorescein (FAM-PEI synthesis), secondary amine assay and transfection evaluation are described in Supplementary Methods.

### Cloning of SpCas9-NLS-mono avidin

SpCas9 constructs with a C-terminal 17 aa linker followed by monoavidin were cloned by an overlap extension PCR. Primers are listed in Table S2. Fragment 1 was amplified using an internal EcoRI site in pMJ915 (Lin *et al*, 2014) and primers SpCas9_EcoRI_445_for and NLS_18linker_rev using iProof High-Fidelity DNA Polymerase (Biorad, USA). Fragment 2 was amplified from pRSET-mSA encoding monoavidin for high affinity binding of one biotin (Lim *et al*, 2013) using primers 18linker_avidin_for and avidin_NotI_rev, and VENT DNA polymerase (NEB, USA). Fragments 1 and 2 were combined and subjected to 10 x overlap extension, before primers SpCas9_EcoRI_445_for and avidin_NotI_rev were added for amplification. The resulting fragment was digested with EcoRI and NotI, and ligated into pMJ806 (Jinek *et al*, 2012) with Kanamycin resistance. pMJ915 and pMJ806 were gifts from Jennifer Doudna (Addgene plasmid # 69090 and #39312) and pRSET-mSA was a gift from Sheldon Park (Addgene plasmid # 39860).

The resulting fusion construct contained an N-terminal hexahistidine-maltose binding protein (His-MBP) tag, followed by a tobacco etch virus (TEV) protease cleavage site and wild type SpCas9 with two C-terminal SV40 nuclear localization signals, and lastly an 17 aa linker to monoavidin. Plasmid Map of Cas9MAV included in Supplementary Figure S1

### Purification of Cas9 proteins

SpCas9 fusion constructs were expressed and purified essentially as described previously (Jinek *et al*, 2012). Briefly, proteins were expressed in BL21(DE3) Rosetta2 cells grown in LB media at 18°C for 16 h following induction with 0.2 mM IPTG at OD_600_ = 0.8. The cell pellet was lysed in 500 mM NaCl, 5 mM imidazole, 20 mM Tris-HCl pH 8, 1 mM PMSF and 2 mM B-me, and disrupted by sonication. The cleared lysate was subjected to Ni affinity chromatography using two prepacked 5 mL HisTrap columns/1-2 L cell culture. The columns were extensively washed first in 20 mM Tris pH 8.0, 250 mM NaCl, 5 mM imidazole pH 8.0, 2 mM B-me and after, in 20 mM HEPES pH 7.5, 200 mM KCl, 10 % glycerol, 0.5 mM DTT, before elution with 250 mM imidazole. The His-MBP affinity tag was removed by overnight TEV protease cleavage w/o dialysis.

The cleaved Cas9 protein was separated from the tag and co-purifying nucleic acids by purification on a 5 mL Heparin HiTrap column eluting with a linear gradient from 200 mM - 1 M KCl over 12 CV. Lastly, a gel filtration on Superdex 200 Increase 10/300 GL in 5% glycerol, 250 mM KCl, 20 mM HEPES pH 7.5 separated the nucleic-acid bound protein from the clean SpCas9 protein. Eluted protein was concentrated to ~10 mg/mL, flash-frozen in liquid nitrogen and stored at −80°C.

### RNP Formation

2.1µl of 1x phosphate buffered saline (sterile and 0.22um filtered), 1.7µl of Cas9 (11-13mg/ml) or Cas9MAV, 2.1µl of 30μM to 200μM sgRNA (concentration varied with respect to Cas9 molarity to maintain 1:1 ratio) were combined in a sterile PCR tube, vortexed gently and incubated for 20 minutes at 25°C. For HDR experiments 5-20µl of 300ng/µl biotinylated or non-biotinylated donors are added at 15 minutes into incubation, with the concentration added adapted to concentration of Cas9MAV-RNP used to maintain a complete binding of MAV within donor. Biotin:monoavidin association occurs within 5 minutes.

### Donor DNA

Donor DNA was created by PCR amplification from gblocks (CXCR4 and EMX1 loci) and for TLR donor from a plasmid kindly donated by Francis Robert from the Pelletier Lab (Department of Biochemistry, McGill). Details of primers and gblocks are found in Supplementary Table S2. For biotinylated donors, substitution of forward or reverse primers with 5’ biotinylated modifications was performed. All oligos were quantified after PCR using nanodrop and concentration of working stocks were 300 ng/µl in all cases. Oligos were used without any further purification. Oligonucleotides used in this research were purchased from IDT (Integrated DNA technologies, USA) and BioCorp (Montreal, Quebec).

### PEI:RNP Polyplex Formation

After formation of RNP, 30µl of PEI (25 kDa 1mg/ml) was added to the RNP and vortexed. The resulting solution was incubated for 20 minutes at 25°C. After incubation 71.5 µl of DMEM was added and tubes incubated at 37°C for 20 minutes to temperature equilibrate with cells to be transfected. Polyplex solution was added dropwise to cells. The above protocol functions for transfecting one well of a 6 well plate, or three wells of a 12 well plate. In the case of larger transfections, quantities were scaled by the estimated cell density of culture vessel. In the case of 6 well plates we estimate 1 million cells at point of transfection.

### SEC-MALS

A size-exclusion chromatography multi-angle light scattering (SEC-MALS) experiment for Cas9MAV was prepared by injecting 50µl sample at 3 mg/ml onto a Superose 6 increase 10/300 GL column (Sigma) equilibrated in 20 mM HEPES, pH 7.5 and 150 mM KCl at a flow rate of 0.3 ml/min. The eluted peak was analyzed using a Wyatt miniDAWN TREOS multi-angle light scattering instrument and a Wyatt Optilab rEX differential refractometer. Data were evaluated using the ASTRA 5.3.4 software with BSA at 5 mg/ml as a reference.

### Dynamic Light Scattering (DLS)

Cas9MAV/sgRNA/dDNA complex hydrodynamic radius and PEI polyplex dimensions were evaluated using a Malvern Nanosizer S dynamic light scattering instrument. Measurements were performed with biotinylated donor DNA concentrations of 2 uM, and preformed Cas9/sgRNA concentrations were adjusted accordingly in a 1:1 molar ratio. All samples were measured in the same buffer (150 mM KCL, 20 mM HEPES pH 7.5) at 20 µl in a Zen2112 cuvette. The flow cell was equilibrated to 37°C to mirror the conditions of cell culture that polyplexes and Cas9 variant would exist during transfection. Measurements were performed for 1 min with a minimum of 15 measurements. PEI conc was 1mg/ml.

### DNA extraction

Cells were harvested using 0.025% trypsin (0.5 ml per well for 12 well and 1 ml for 6 well plates) after a 1xPBS wash and centrifuged in 1.5ml eppendorfs at 8000k for 5 minutes. Supernatant was discarded and pellet washed with 0.5ml PBS. Cells were spun again and supernatant discarded. Full protocol for extraction is detailed in Supplementary Methods. Homemade lysis, column binding, wash and elution buffers compositions are in Supplementary Methods. Proteinase K and RNase 1 were sourced from Sigma Aldrich.

### RNA extraction

For cellular RNA Trizol extraction following manufacturer’s protocol (Thermofisher, USA) and Direct-zol clean up columns (Zymo Research) were used for preparation of RNA for RT-PCR validation of genomic edits. DNAse1 treatment for enzymatic degradation of genomic DNA was performed following manufacturer’s protocol. DNA degradation was verified by gel electrophoresis. DNase1 and buffer was supplied by NEB, USA.

### PCR

Genomic PCR was performed with either Q5 2X master mix or Phusion polymerase (NEB, USA). PCR was used for donor DNA creation using IDT gBlocks as templates. All oligos used are specified in Suppplemental Table S2.

### RT-PCR

Reverse transcription was performed using the High reverse transcriptase kit (Biobasic, Canada). For PCR step 2 microliters of solution were used in the standard Phusion master mix protocol (NEB, USA).

### Gel Electrophoresis

For analysis of PCR products, and enzymatic digests, 2% agarose (Biobasic, Montreal) gels were prepared and a Fluoro-loading dye (Zmtech Montreal) was used to visualize the DNA (2µl per 25µl volume). Gels were run at 90 volts in 1x TAE buffer. For RNA analysis urea polyacrylamide gels (12%) were used and run at 120V for one hour with a pre-run prior to loading of half an hour at 130V in 1X TBE buffer. RNA gels were stained with a Ethidium bromide staining solution (10µl EtBr 10mg/ml in 30ml TBE). Gel images were recorded using UV illumination and imaging gel dock.

### T7 Endonuclease Indel assay

T7 Indel assay was performed according to the protocol of Guschin et al (Guschin *et al*, 2010) with products being quantified by agarose gel electrophoresis. PCR amplification of genomic edited and unedited DNA was performed using Q5 polymerase. All T7 endonuclease assay samples were performed with minimum of 3 biological replicates. ImageJ was used for analysis and quantification of band intensity detailed in Supplementary Methods. Results were analyzed by the ANOVA and Tukey statistical method using the R programming language { RAlanguageanden:uc}). The ggplot (ISBN 9780387981406) package in R was used to generate the graphs.

### Restriction Digest HDR assay

HDR frequency was assessed by restriction digest analysis. Genomic loci chosen were analysed by NEBcutter 2.0 to identify any restriction digest sequences in endogenous DNA. Excluding those enzymes with palindromic sequences present, specific enzymes without palindrome sequences were used to create a donor inserts. In the case of CXCR4 and EMX1, the donor sequence TAG-HindIII and BamHI (respectively, see Figure 2 and 3) were inserted into gblocks with homology arms matching the genomic loci. Details of gblock sequence and subsequent PCR method for creating donors of various lengths are included in the Supplementary Data. Restriction digest was performed after amplification from DNA extracted from edited and unedited cells following the manufacturer’s protocol (NEB) for cut smart High fidelity restriction enzymes. All HDR restriction digest assays were performed with a minimum of 3 biological replicates. In brief, 5 µl of PCR amplicon was diluted in 35 µl nuclease free water and 5 µl cut smart buffer. 1µl of high fidelity restriction enzyme (HindIII or BamHI) was added and vortex, volume was adjusted to a final volume of 50 µl. Reaction was incubated at 37°C for 15 minutes and quenched with Fluoro-DNA loading dye 6X (Zmtech Scientific, Canada), before analysis directly on 2% agarose gels. Gel images were analysed in ImageJ and percentage HDR was calculated according to equation 2 (Supplementary Methods). Results were analyzed by the ANOVA and Tukey statistical method using the R programming language { RAlanguageanden:uc}. The ggplot (ISBN 9780387981406) package in R was used to generate the graphs.

### TLR HDR assay

TLR assay was performed using cells transfected by AAV virus with pCVL-TL-dsRed Reporter 2.1 (VF2468 ZFN target) EF1a inserted at a safe harbour loci (Robert *et al*, 2015). The TLR transfected cell line was a kind gift from Francis Robert in the Pelletier lab. Cells were plated into 12 well plates and split into 6 well plates 24 hours after transfection. Cell were then grown for 5 days with samples. Control Cas9 plasmid transfections were performed by PEI transfection (1ug plasmid DNA, 30µl PEI and 300 µl DMEM for a 2 well transfection). PEI:Cas9MAV was transfected into cells with 5’ b donor, 3’ b donor and 5’3’ b donor. Cells were prepared for flow cytometry at 48 hrs and 96 hrs. Further details in Supplementary Methods.

The Cas9 plasmid was a gift from Jerry Pelletier (McGill University).

### Fluorescent Microscopy of TLR assay Cells

For TLR assay cell plates (12 well and 6 well) GFP and RFP fluorescent images were collected using a Zeiss AX10 inverted fluorescence microscope and standard GFP and RFP filters. Image acquisition required a AxioCam MRm camera and Axiovision software. TIFF images were processed in ImageJ.

### Cloning and Sanger sequencing

PCR products for sanger sequencing were cloned into bacterial blunt end cloning vectors (pCR, pUCM) as part of blunt end cloning kits (Thermo Fisher ZeroBlunt kit and Biobasic pUCM-T cloning vector kit). Plasmids were transformed into XL1 Blue competent cells. Cells were grown on selective plates and colonies picked for overnight broth culture and subsequent mini-prepping (kit). Where pUCM-T vector was used a 1-TAQ (NEB, USA) polymerase was used to give the adenosine overhang. Sanger sequencing was performed by the McGill University and Génome Québec Innovation Centre (Montreal, Quebec). Sequence analysis for Sanger sequencing was performed in CLC Main Workbench (QIAGEN Bioinformatics, Redwood City, CA, USA) for sequence alignment and identification of donor insert sequences.

### Ion Torrent

Ion Torrent sequencing of DNA samples was performed on a ThermoFisher Scientific Ion Torrent Personal Genome Machine ^TM^ (PGM) apparatus using Ion 316 chip kit v2 (ThermoFisher Scientific, 4483324), with potentially 6 million reads. Samples were multiplexed upon the 316 chip by barcoding of 9 samples by PCR prior to the Ion Torrent work flow. For the second round of sequencing focusing on the 137 bp donor, extracted DNA samples were amplified using genomic primers generating a 800bp product from DNA outside of the insert sequence, and EMX1 was amplified to produce a 600bp product accounting and excluding for any residual donor. A subsequent amplification was performed with ion torrent barcoded primers for deconvolution of reads. Experimental conditions were tested in quadruplicate or triplicate. Details of barcoding primers is included in Supplementary Tables S4, S6. After barcoding PCR, samples were run on a 2% gel and each band pertaining to the library amplicons (approximately 200bp) were excised and gel extraction performed (Qiagen gel extraction kit). DNA was quantified by Qubit (picogreen) and diluted to 8 pmol prior to Ion Torrent work flow.

### Ion Torrent CRISPR/Cas9 read processing

Sequenced products were filtered for a minimum length of 61 nucleotides (minimum expected distance to target subsequences of interest – described below) and mapped to a custom BLAST database (v2.5.0) consisting of 14 sequences (a combination of seven barcodes [plus primers] and two expected sequences [no insertion or the insertion of a stop codon and HindIII restriction enzyme site]). BLAST mapped libraries were then analyzed for two target subsequences: 1) the presence of an insertion (TAGAAGCTT) or 2) absence of any change in the canonical sequence near the CRISPR/Cas9 insertion site (CCAGGAT). Aligned reads that did not completely overlap with the target subsequences were removed (‘truncated’ reads). Any alignments that resembled a partial insertion (e.g., TAGA, TAGAA, TAG-A, etc.) or non-insertion (based on BLAST alignments) were then analyzed for their frequency of insertions or deletions (indels) at a given position of the target site, including five nucleotides up-/down-stream of the target subsequence (see Supplementary Data). A total of 793,117 sequenced CRISPR/Cas9 products were produced, where 2.522%, 0.104%, and 97.374% of products were too short to be aligned to expected product sequences, unmappable by the BLAST algorithm, or able to be aligned by BLAST to the expected product sequences, respectively. Of the 772,286 mappable products: 728,664 (91.873% of total sequenced products) contained a perfect match to the target subsequences; 16,083 (2.028%) produced alignments that did not overlap with expected target subsequences; and the remaining 27,539 (3.472%) contained partial insertions/non-insertions.

## Acknowledgements

We would like to thank Francis Robert and Jerry Pelletier (helpful discussions, loan of equipment, and reagents), Bozena Samborska from Russell Jones’ lab (gift of reagents and cell lines), Shawn McGuirk from Julie St. Pierre’s lab, Jutta Steinberger from Jerry Pelletier’s lab, Sidong Huang (material gifts of cell lines), Alba Guarne and Kalle Gehring (providing access to DLS and SEC-MALS, respectively).

The authors would like to acknowledge the financial support of the Faculty of Medicine, McGill University (Akavia), Canadian Institutes of Health Research (CIHR grant MOP-133535 to Nagar; CIHR grant MOP-142451 to Dostie), and the Natural Sciences and Engineering Research Council of Canada (NSERC Discovery grant to Blanchette).

## Author Contributions

PJ Roche: Conceptualization, Investigation, Methodology, Project Administration, Supervision, Visualization, Writing - original draft and Writing - review & editing

H Gytz: Conceptualization, Investigation, Methodology, Resources, Visualization, Writing - original draft and Writing - review & editing

F Hussain: Formal Analysis, Investigation, Visualization and Writing - original draft CJF Cameron: Data Curation, Formal Analysis, Software, Validation and Visualization M Blanchette: Software, Funding Acquisition and Supervision

D Paquette: Resources, Validation

J Dostie: Funding Acquisition, Resources and Validation

B Nagar: Funding Acquisition, Supervision, Resources and Writing - review & editing

UD Akavia: Funding Acquisition, Conceptualization, Project Administration, Supervision, Visualization, Writing - original draft and Writing - review & editing

## Conflict of Interest

The authors have no competing financial interests.

## Supplementary Figure Legends

<u>Figure S1</u> Plasmid map of Cas9MAV

<u>Figure S2</u> CXCR4 HindIII digests after cell growth

CXCR4 HindIII digests were run on an agarose gel at 115V for 20 minutes. For both uncut and cut: DNA ladder (lane 1, 1000bp), CF1b (lanes 2, 6, 10), Fas1b (lanes 3, 7, 11), Ras1b (4, 8, 12), CF3b (5, 9, 13). In the HindIII gel, lane 14 includes uncut CF1b as a control.

<u>Figure S3</u> Indel frequency at CXCR4 and EMX1 loci

Comparison of indels formed with nucleofection of Wild type Cas9NLS and Cas9MAV RNPs

<u>Figure S4</u> Comparison of Cell viability with each of the transfection methodologies

<u>Figure S5</u> Traffic Light report assay for HDR and NHEJ

Data was assessed at 48hr (**A**) and 96hrs (**B**) post transfection with PEI:Cas9MAV and the Cas9NLS plasmid. Data is presented using the quadrant gating approach used previously by Certo et al. Control Sample quadrant values used to normalize each time period and set the gating to exclude dead cells. Cells in lower left quadrant represent the unedited population. Cells in upper left quadrant express GFP, hence HDR. Those cells in upper right are co-expressing GFP and RFP, are both HDR and NHEJ. Cells found in the lower right quadrant are expressing RFP representing NHEJ.

Each experiment was repeated twice for flow analysis apart from combined 5’3’ donor. 5’ in the context of this experiment means Forward primer biotin generated donor. 3’ means reverse primer with modified biotin. 5’3’b refers to a donor generated as a combination of both biotinylated primers. Sample 3’b at 96hrs had few to no surviving cells post harvesting by trypsin and PBS washes.

## Supplementary Materials and Methods

### CXCR4 Re-growth experiments

An important hypothesis considered was if cells are edited at high frequency (99%) then the edit should persist in cell population as the most numerous sequence. If the edit is lower frequency after a period of re-growth, edited cells are regularly out competed and this presents a significant challenge to single cell cloning for formation of stable cell lines. If the results of the CXCR4 experiment as confirmed in figure 3 at genomic and mRNA levels are consistent, the edit should persist after cell passaging and aggressive dilution of total cell number for the seeding of subsequent plates.

Cells were edited at CXCR4 using PEI-Cas9 system with biotinylated donors as per standard protocol and cell density. After 48hrs they were split 10:1 into a 6 well plate. Hypothesis is that if edit is 100% or near, the edit will still persist after aggressive dilution and 7 days re-growth, validating that the edit is stable and inheritable. Figure S2, confirms the edit persisted after a 10:1 passage into 6 cell plates and 7 days re-growth. The experiment confirms that all donors induce persistent, high level HDR at this loci.

### Nucleofection Transfection of the Cas9MAV

Transfection of the Cas9MAV RNP was evaluated by the performance on Amaxa Nucleofector II (Lonza), using the T7 indel assay as a means to confirm nuclease performance.

Briefly HEK-293T cells were transfected with the Amaxa Nucleofector^®^ Kit V (Program Q-001) with a GFP positive control performed to confirm nucleofection instrument functioned. Cells were harvested and counted. 1 million cells were resuspended in Kit V buffer with RNP prepared as per RNP transfection method.

The indel frequency was assessed as near equivalent to PEI transfection, for both Wild type Cas9 (no NLS) and Cas9MAV at both loci. In addition it confirms the Cas9MAV system can be introduced by alternative transfection systems.

### Comparison of Cell Viability when transfecting Cas9MAV

Cell viability of transfection method was considered an important factor in the choice of PEI as the means of introducing Cas9MAV to the cells. Maintaining a high cell viability (greater than 80%) means a great number of cells can be potentially edited. Figure S4 details the percentage live cells assessed by trypan blue staining after 48hrs. Three methods are compared, PEI, Lipofectamine 3000 and Amaxa nucleofection using kit V for the transfection of Cas9MAV in HEK293T cells. None of the methods in our hands using the protocols detailed significantly affecting cell viability.

### TLR Flow analysis

The second locus we investigated is a variant of the Traffic light reporter assay. The cell line was a gift from Prof Jerry Pelletier and Francis Robert (Biochemistry, McGill University). Donor template was provided on a plasmid backbone and amplified using TLR donor F and R primers for biotinylation (see oligo table in Supplementary Table S2). Cells were transfected using PEI:Cas9MAV and biotinylated donor DNA to induce GFP expression in the case of HDR, or mCherry if only indels are formed via NHEJ.

Data is presented using the quadrant gating approach used previously by Certo et al (Certo *et al*, 2011) in Supplementary Figure S5. Control Sample quadrant values used to normalise each time period and set the gating to exclude dead cells. Cells in lower left quadrant represent the unedited population. Each experiment was repeated twice for flow analysis apart from combined 5’3’ donor. 5’ in the context of this experiment means Forward primer biotin generated donor. 3’ means reverse primer with modified biotin. 5’3’b refers to a donor generated as a combination of both biotinylated primers.

For each sample 2000-5000 live cells were gated and analysed for RFP and GFP expression using the Guava flow cytometer. Sample 3’b at 96hrs had few to no surviving cells post harvesting by trypsin and PBS washes.

Cells in upper left quadrant express GFP, hence HDR. Those cells in upper right are co-expressing GFP and RFP, are both HDR and NHEJ. Cells found in the lower right quadrant are expressing RFP, representing NHEJ.

Cas9 plasmid control is included to give a measure of the standard performance of plasmid vectors expressing Cas9, but also the NHEJ frequency normally observed in the absence of donor. It should be noted Cas9 plasmid requires 4-7 days to produce a reliable signal and only on the 4th day did the plasmid system record a indicative signal and hence it’s inclusion only in the 96hr data set. Overall performance of Cas9MAV was assessed by averaging the performance of all donors (n=4 repeats, 2 for each donor type) and for each time period (48hrs and 96hrs).

### Suppression of NHEJ using PEI:Cas9MAV RNP system

Suppression of NHEJ was observed where HDR frequency was found to be in high order by HINDIII restriction site insertion into genomic loci (CXCR4). Figure S6 demonstrates an example where in combination with the RNP system, Nocodazole was administered to investigate the effects of cell cycle arrest. Principally the effect was negligible, as HDR rates were statistically irrelevant from using the RNP system without cell cycle arrest. Therefore, nocodazole was not used in further experiments.

The top bands in lanes 2-9 correspond to the uncut sample, whereas the bottom bands are the cleaved products, and slightly lower are the primer-dimers. As seen in the gel, different donors led to different amounts of NHEJ. For example in lane 2, it is seen that nocodazole treated samples with the CF1 donor produced no NHEJ (no bottom band is visible) whereas in the zoomed lanes 6 and 7 it is clear that T7 produced a cut with the Fas2 donor. Technical replicates were performed and further analysis was done in ImageJ to determine the extent of NHEJ with all the variants. Using the intensities of the bands, the following equation was used to calculated indel frequency:

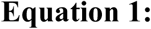

### Assessing HDR frequency by insertion of restriction digest palindromic sequences

As a rapid means of assessing HDR frequency EMX1 and CXCR4 donors were designed with a restriction digest site not present in the genomic sequence. The loci were assessed using NEB cutter restriction digest predictor before selection of enzymes was performed. In the case of EMX1 BamH1 was chosen and CXCR4 HindIII was used. Once HDR was completed after 48hrs comparison of samples against an unedited control were performed. An example of the results and analysis method are shown below:

The top bands in lanes 2-10 correspond to the uncut sample, whereas the middle two bands are the cleaved products, and slightly lower are the primer-dimers. Similar to the results from the T7 tests, the various donors led to different amounts of HDR. For example in lane 2, it is seen that Cas9NLS samples with the CF1 donor produced an apparent 100% HDR (no top band is visible) whereas in the lanes 3 and 4 (Cas9NLS with Fas1 and Fas2) HindIII did not completely cleave the upper band (so there is less than 100% HDR). Both biological and technical replicates were performed and ImageJ was used to determine the HDR efficiency with all the variants.

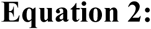

### Deeper Evaluation of HDR by Ion Torrent Sequencing

A key facet of investigating the high frequency of HDR was to explore the direct sequence of libraries derived from our CXCR4 HDR experiments (fig.2 main paper). A criticism of HDR assay by restriction digest insertion is that it can under and overestimate HDR based on band intensity, blurring efficiency of cut and experimental error. While best experimental practice can be followed and the approach has been applied widely by Doudna et al (Lin *et al*, 2014) among others. In brief the introduction of a palindromic sequence enables rapid identification of edits. The degree to which the PCR product from genomic DNA is cleaved based on the introduction gives a measure of the HDR events. Alternative approaches such as reporter assays, can be open to other variables such as donor length and whether the insert is inframe and transcribed correctly. To delve into the occurrence of HDR in this manuscript, Ion torrent sequencing was chosen. By using next generation sequencing, we can explore frequencies for complete HDR inserts, partial inserts/indels and un-edited sequences. With greater coverage than Sanger, which is biased towards the most dominant sequences, a more accurate evaluation of the HDR frequency than the gels was attempted.

## Supplementaary Methods

### Genomic PCR

All primers were ordered from either IDT or Biocorp and diluted in Rnase free water to 100uM stocks for storage at −20°C. Working stocks of 10uM for all primers were prepared and stored at - 20°C until required. PCR was performed with Phusion, Q5 and One Taq (NEB) in 25 µl volumes according to the manufacturer’s instructions.

### PCR generation of DNA donors and Biotinylated donors

Generation of donor DNA was performed from gBLOCKS design with restriction digest sites specific to the gene we hoped to edit. All donors of different lengths were performed using match forward and reverse primers to generate asymmetric, equal homology arms variants and biotinylated versions created by substitution of either reverse or forward primer as per diagram in figure 3. Dependent upon the quality of PCR band, gel extraction was performed for clean up.

Donor design was based on a series of conclusions from papers listed in Supplementary Table S1. CF1 and CF3 were designed with 600 and 400bp homology arms that were seen to be optimal for larger double stranded donors. For Fas1 and Fas2, a short sub-optimal double stranded donors. We applied asymmetric HA based on the 5’ 37bp arm being shorter than the distal 93bp arm after the insert. Insertion was design to break the PAM and part of the guide sequence to disrupt re-cutting that could mitigate HDR identification. This design rule was incorporated in to CXCR4 and EMX1 donor design templates.

We opted for gBLOCK design templates and PCR reaction of donors, based on ease of experimentation and speed. It also allowed the flexibility to mix 5’ biotinylated primers into the reactions as substitutes to create biotinylated versions rapidly.

Double stranded donors were applied in all cases, deliberately for positive reason (CF1/CF3) and negative reasons (Fas1/Fas2) and the choice was borne out in the relatively poor performance of shorter donors when non-biotinylated versions were applied in fig.3. In choosing biotinylated versions we opted for CF1b, F denotes the forward primer carries the biotin, and this was the case with Fas1b and CF3b but Ras1b applied a reverse primer modification.

For donors with TLR, we used a 111 base pair sequence designed by Francis Robert of the Pelletier lab. In a different approach we used a double stranded donor using with 5’ or 3’ biotinylation, or dual biotinylation, to study the impact. 5’ and 3’ labelling denotes the orientation of the original single stranded donor design. The 3’ mod implies the location of the biotin modification on the antisense strand, as opposed to the 5’ label which is upon the sense strand. The donor design is asymmetric and is shorter than the recommended 137 bp used by Richardson et al (Richardson *et al*, 2016).

For EMX1 we deliberately mitigated the HA to below 200bp (total donor length 400bp plus tag repeat sequence and BamH1 site). The intention was to evaluate whether another sub-optimal design as was performed with TLR, could be recovered by using the PEI transfection approach. In achieving ~70% HDR, it was indicative that PEI nuclear location can rescue or mitigate the conventional donor design rules. We have tested the biotinylated version of the EMX1 donor, and found it followed the performance trend of the Fas1b and Ras2b dDNA, recovering a higher HDR frequency by benefiting from donor co-localisation to DSB.

An aspect unexplored in the mechanistic sense in this work is the impact of removing diffusion and nuclear translocation from the dDNA by attachment to Cas9MAV. With other transfection methodologies, nucleofection excepted, the challenges of nuclear import remain and it is possible less than 30% reaches the nucleus from studies of plasmids and presuming free donor DNA diffusion occurs slowly (3×10^-8^cm/s) (Zhou *et al*, 2004), unless under the influence of active transport and importins. This consideration lead us to evaluate the importance of donor:RNP co-localisation and concentration. In future work we hope to evaluate accurate quantities of Cas9MAV:donor conjugate within the nucleus.

### sgRNA synthesis

IVT one piece is made from attaching T7 promoter upstream of guide sequence to be tested, and a overlap sequence. Overlap sequence matches the reverse complement of the S.pyg scaffold.

### Example for CXCR4

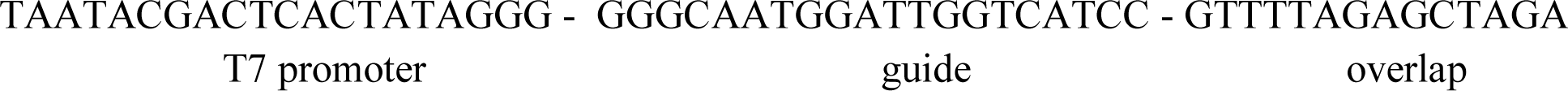

Protocol follows the NEB Engen protocol, with one minor alteration. All sgRNA were prepared by Engen sgRNA synthesis by the appending of a T7 promoter sequence and 3’ hybridisation sequence to the reverse complement strand. Using the combination of bacterial polymerases and T7 transcriptase in the kit, DNA strand elongation occurs to create a double strand DNA template for the T7. Differing from the standard NEB protocol, reactions were carried out over 12-18hrs, after which DNAse1 treatment was performed for 30mins. DNase1 was heat deactivated in the presence of EDTA to remove MgCl2. sgRNA can be cleaned up using a rapid RNeasy spin columns or similar kits such as RNA clean and concentrator (Zymo Research USA). sgRNA was confirmed for molecular weight using 6M Urea polyacylamide gel electrophoresis. RNA was then quantified using a nanodrop and molarity calculated. Working stocks of sgRNA were prepared at 300-1000uM, with addition of RNase inhibitors and kept at - 20°C. Storage solutions were kept at −80°C with Rnase inhibitor at manufacturer’s recommende concentration.

### DNA extraction methodology

Pellet was resuspended in 0.2ml PBS, 0.2 ml Lysis buffer with 20mg/ml Proteinase K and incubated for 2-12hrs at 54°C on a dry heating block. After, samples were incubated with 2µl RNAse (10mg/ml) for 30 minutes. 0.2ml 4M GuHCl binding buffer and 0.2ml EtOH was added and the solution inclusive of precipitation was added to econo-spin silica columns. The samples were centrifuged for 3 mins (8000K) and supernatant discarded. 500µl of wash buffer was added to column, which was spun (3min, 8000K). The wash step was repeated again and supernatant discarded. Column was supn for 1 minute to dry silica membrane of column and replace collection tube with an 1.5ml eppendorf tube. Sample was eluted with 50-100µl of elution buffer or DEPC water by centrifugation (1 minute, 8000K). Quantification of extracted DNA was performed using a Nanodrop spectrometer.

#### Buffer compositions

> Lysis: 10mM Tris 2mM EDTA 1% SDS
>
> Binding buffer: 3M GuHCl 3.75M NH_4_Ac pH 6
>
> Wash buffer: 10 mM Tris-HCl pH 7.5, 80% ethanol
>
> Elution: MilliQ water or 10mM Tris:EDTA buffer at pH 8

### PEI preparation

The degree of protonation of PEI has an impact upon polyplex formation and transfection efficiency. We applied a ninhydrin assay to assess the protonation of secondary amine. At pH 7.4 it is suggested that the ideal 40-50% protonation occurs and using the assay an absorbance of 0.025 to 0.03 for PEI incubated with ninhydrin is considered the goal of PEI solution preparation.

In brief a 100mM Ninhydrin solution was prepared in milliQ water. 333µl of Ninhydrin stock and 666µl of PEI solution were vortexed and heated to 75°C for 20 minutes in a dry bath, followed by quenching on ice for 5 minutes. Solution was measured by UV/VIS (487nm) and the absorbance recorded. For evaluation of PEI consistency, a calibration curve was prepared with PEI solutions adjusted from pH1 to pH 14. 40-50% protonation of secondary amines equates to pH7-7.4.

### PEI labelling

Adapting the method of Godbey et al (Godbey *et al*, 1999), FAM-nhs ester was substituted for oregon green. Briefly PEI (10mg/ml, supplier) was dissolved in 0.1M Sodium bicarbonate and reaction of the primary/secondary amines with FAM-NHS ester (50µl per ml 10mg/ml supplier) was performed at standard laboratory conditions (1atm, 25°C) and magnetically stirred. Vessel was enclosed to exclude light and photobleaching of fluorescent dye. Resulting FAM-labelled PEI was aliquoted into 200 µl volumes and stored at −20°C until required. For Fam labelling of polyplexes, aliquots are defrosted and vortexed, before adjustment to room temperature. 2-5µl of total 30µl (10mg/ml) unlabelled PEI used to form RNP polyplexes is substituted.

### Transfection with FAM-labelled PEI:Cas9MAV

Polyplexes were formed using the standard PEI:Cas9MAV protocol (Supplementary Methods) substitute 5µl of unlabelled PEI for FAM:PEI. HEK293T were seeded into a 6 well plate 24hrs before to adhere (seeding density 3×10^5^) to achieve 60-70% confluence at point of transfection. After 24hrs cells were washed with 1x PBS and media refreshed before fluorescent image acquisition using an EVOS cell imager. ImageJ was used for basic processing and image normalisation.

### Liposomal delivery of Cas9MAV

As a comparative experiment we performed lipofection using Lipofectamine 3000 (LP3K). The basic protocol for lipofection was adapted from Yu et al (Yu *et al*, 2016). To maintain equivalence of experimentation, the same RNP preparation (Cas9MAV, sgRNA concentrations) as for PEI transfection was used for RNP formation and donor addition (biotinylated donors or non-biotinylated donors). 0.3 µl per well of LP3K was used and after ten minutes incubation at 25°C, transfection solutions were added to cells (5.5µl per well). The RNP transfection solution followed the recipe below:

|  |  |
| --- | --- |
| RNP mixture | vol. 5.9µl |
| LP3K | vol. 0.9µl |
| Donor | vol. 5µl |
| OptiMEM | vol. 4.7µl |

On two 12 well plate 3 repeats were created for each non-biotinylated donor (CF1, Fas1, Fas2, CF3) and biotinylated variant (CF1b, Fas1b, Ras1b, CF3b). The other alteration to the PEI transfection protocol reflects LP3K stability in serum rich media. Prior to transfection standard DMEM media was removed and replaced with OptiMEM (Gibco). Two hours after transfection the OptiMEM media was replaced by DMEM. Cells were allowed to grow for 48hrs prior to analysis.

### Ion Torrent Sequencing and Preparation

For ion torrent sequencing design of new primers for a 119 bp region bridging the insert site were designed. Figure S10 shows the genomic location with HDR edit validated by Sanger sequencing. Primer 1 and Primer 2 denote the genomic designed for CXCR4.

In the context of Ion torrent sequencing addition sequences needed to be appended to the forward and reverse primers. At the 5’ end of the forward primer a 10-12 ntp barcoding sequence was added, unique to each sample. The reverse primer was appended at the 5’ end with the P1 key, containing a taqman probe target sequence (CCACTACGCCTCCGCTTTCCTCTCTATGGGCAGTCGGTGAT, R-Ionseq in Supplementary Table S2), as important as part of the ion torrent work flow. Detailed barcodes and library descriptions are included in Supplementary Table S4.

An initial amplification is performed on all samples using One Taq from genomic DNA samples extracted from un-edited and HDR edited cells (different donors, state of biotinylation). Once confirmed by gel electrophoresis (2% agarose), confirming product size as 200 bp and forming a single product. The products for each sample were gel extracted and purified with a Qiagen QIAquick gel extraction kit. Each extraction had between 2-5ng/µl. 5µl of each sample were combined in a clean 1.5ml eppendorf, average sample concentration was given a 4ng/ul and the quantity of total DNA in 35µl was estimated to be 0.14µg, equating to a molarity of 30.46nM. Samples were then combined and diluted to 8 pmol total DNA ready for emulsion PCR as part of the ion torrent work flow. Templating and sequencing were performed according to the Ion torrent ION PGM template OT2 200 kit and ION PGM Hi-Q sequencing protocols.

BC7 library sequencing was not included in data analysis due to minimal sequences completed in comparison to other libraries. Sequences are included in the FastQ data file but not used for further analysis.

Supplementary Table S5 details tables of alignment and frequency of occurrence for each nucleotide that differs from the predicted sequence. Predicted sequences are either the un-edited genomic (length 17bp), or the expected edited sequence (length 19bp). Focusing upon the sequence comparison frequency tables were generated to show where the frequency of alternative base assignments after BLAST analysis.

Attached as Supplementary Dataset 2 (SD2) are the FastQ raw reads, and excel spreadsheet investigating knockin variants by analysis of alternative base pair assignments. The most common knock in variant assessed results from a single nucleotide insertion. Given the nature of Ion torrent sequencing being prone to indel formation this sub-population was assigned separate from complete and accurate knockins, as uncertainty regarding assigning it as a genuine knockin variant was precluded.

Software used to analyze the sequencing results was written in the Python programming language and is attached as Supplementary Dataset SD3.

The second NGS sequencing experiment focused is detailed in Supplementary Table S6, including sequencing subdivisions for inserts (HDR), partial inserts, partial wildtype and wild type sequences, for the CXCR4 locus. Experiments were conducted with PEI or Lipofection with either biotinylated or non-biotinylated 137bp double straned donors. Details of barcodes and primers used as well as corelation between barcode and experimental names used in Figure 4.

### Mycoplasma Testing

For cell culture contamination testing through the period of experimentation (September to December 2017) we applied a PCR based mycoplasma test (Zmtech, Montreal). Cells and culture media were sampled bi weekly and PCR was performed according to manufacturer’s protocol.

## Supplementary Tables

<u>Table S1</u> - Review of Cas9 HDR Improvement strategies

<u>Table S2</u> - Oligonucleotide sequences used for experimentation

<u>Table S3</u> - Summary table for TLR experiments

<u>Table S4</u> - Analysis of Ion torrent sequencing

<u>Table S5</u> - Frequency tables of first sequencing analysis

<u>Table S6</u> - Additional Ion Torrent libraries

## Supplementary Datasets

<u>SD1</u> - PDF file of Sanger sequenced clone raw reads and chromatograms

<u>SD2</u> - FastQ file for the first ion torrent libraries

<u>SD3</u> - Ion torrent sequencing data processing code

<u>SD4</u> - FastQ files for second ion torrent libraries

<u>SD5</u> – Tukey analysis and R-code

---

**Figure S2:**
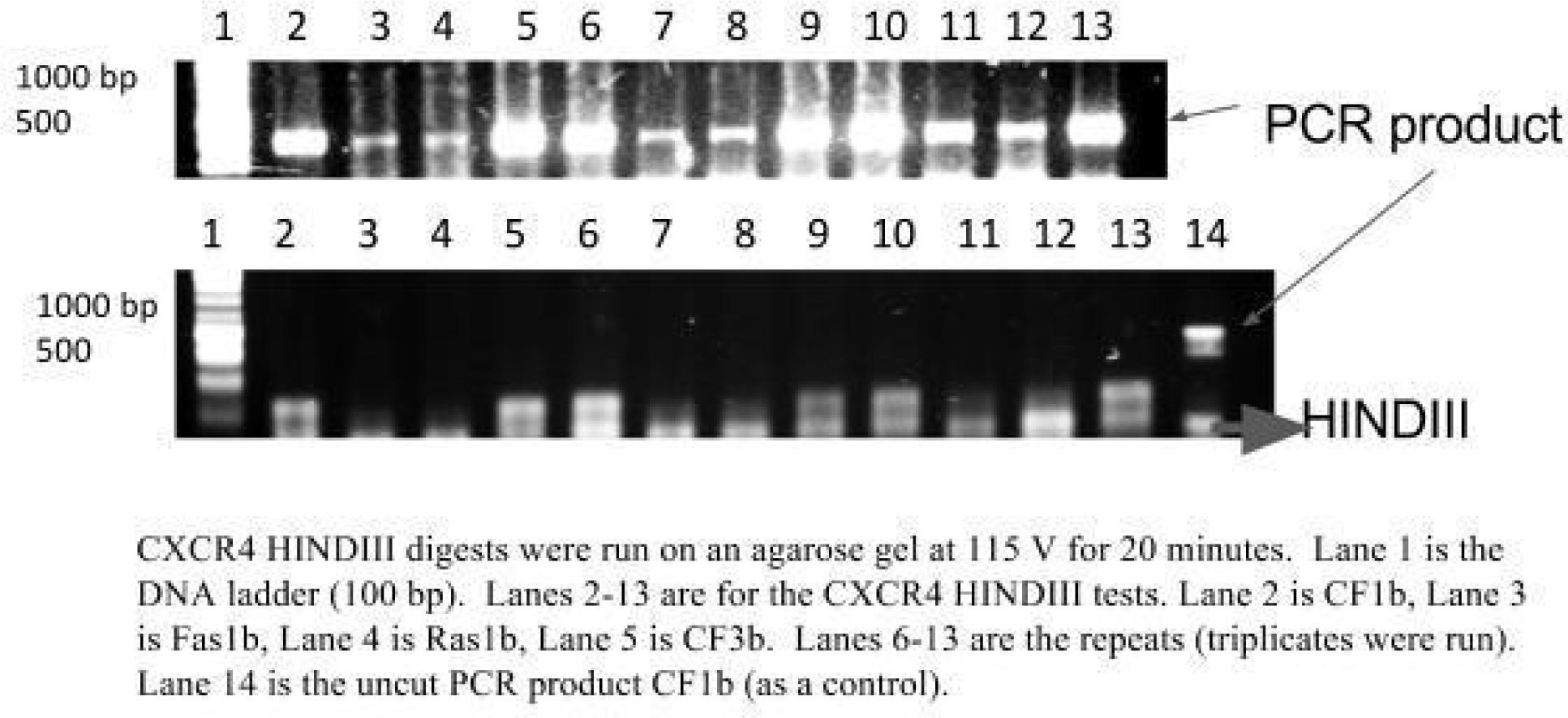
- Growth of HDR edited cells to evaluate the stability of HDR edit after aggressive 10 to 1 dilution of cells.

**Figure S3:**
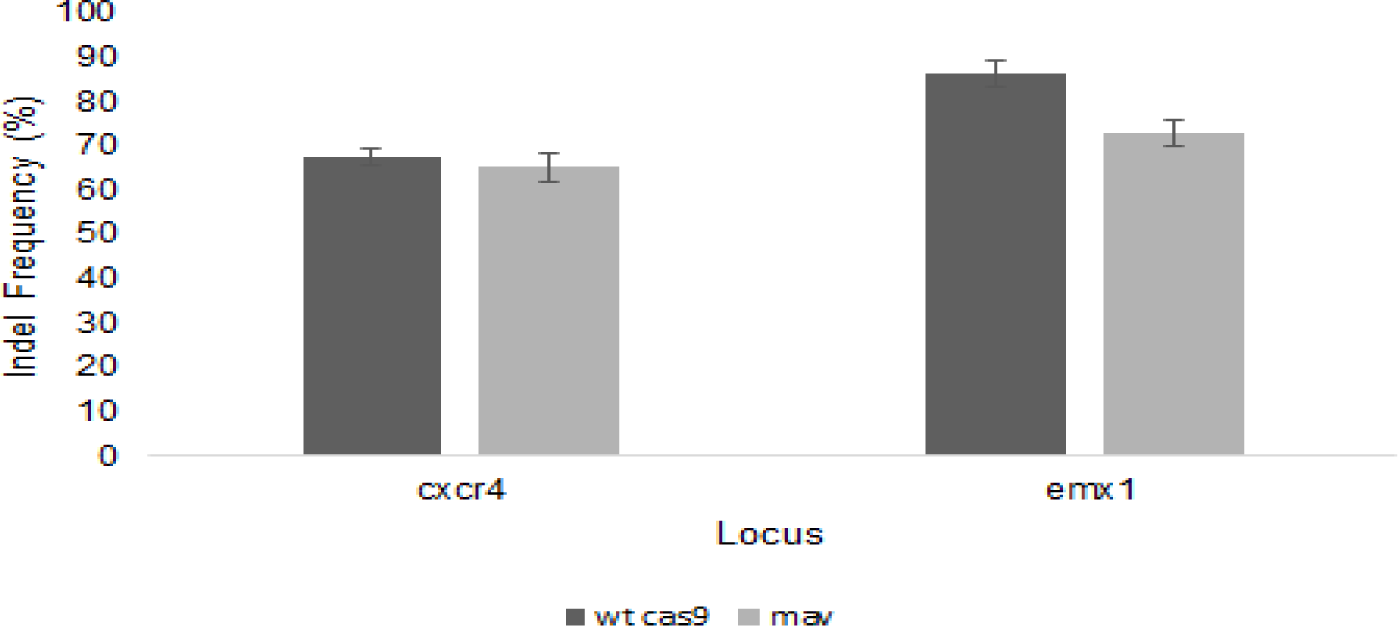
**Indel frequency at CXCR4 and EMX1 loci** nucleofection of Cas9MAV RNP. Comparison of Wild type Cas9NLS and Cas9MAV

**Figure S4:**
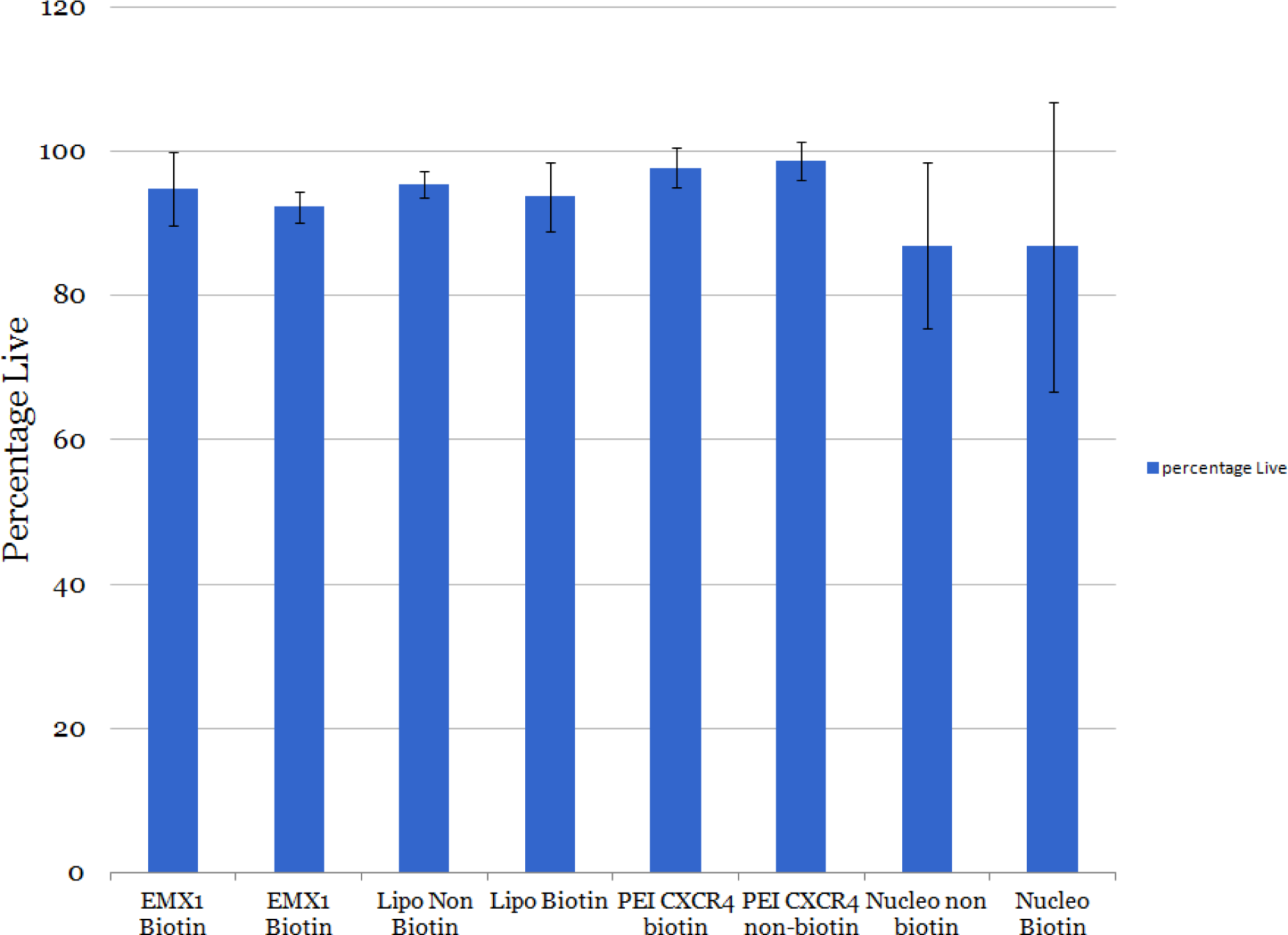
**Comparison of Cell viability with each of the transfection methodologies**

**Figure S6:**
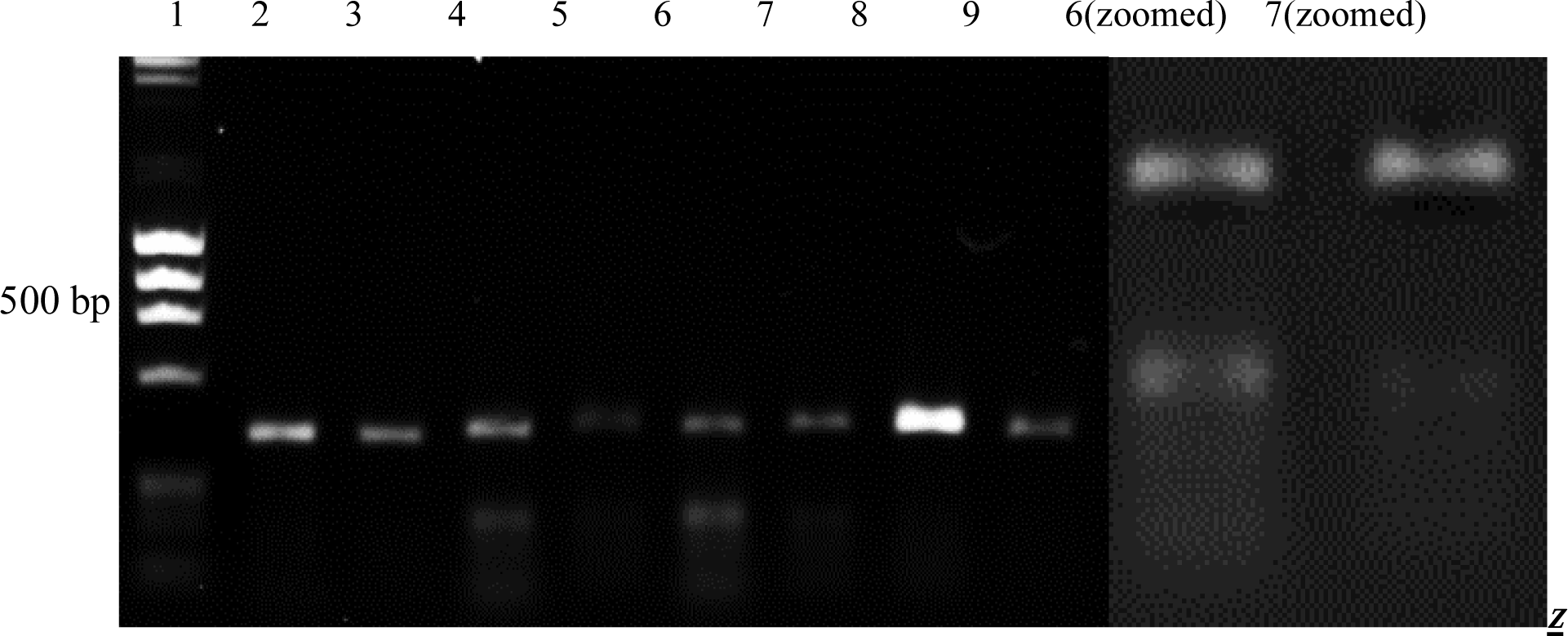
**T7 Endonuclease Assay of Nocodazole Treated Samples from CXCR4 locus**. A 2% TAE agarose gel was run at 90V for 20 minutes. In lane 1, 5 μL of DNA ladder was run. The following lanes contain T7 digest products of the nocodazole treated variants (with unbiotinylated donors) from the CXCR4 locus (5 μL in each lane). The lanes 2-9 show alternating T7 cut products and uncut samples. Lanes 2 and 3 show Cas9-noc using the CF1 donor. Lanes 4 and 5 show Cas9-noc using the Fas1 donor. Lanes 6 and 7 show Cas9-noc using the Fas2 donor. Lanes 8 and 9 show Cas9-noc using the CF3 donor. On the right is a zoomed version of lanes 6 and 7 for easier visualization of the bands.

**Figure S7:**
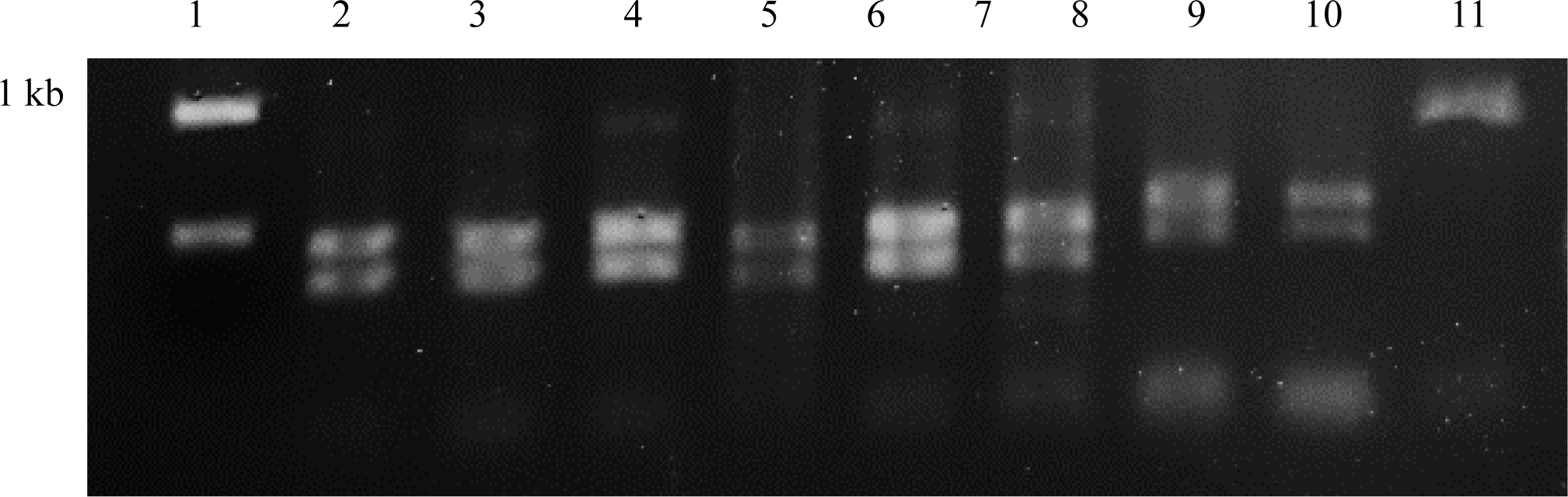
**HindIII Digest Assay of samples from CXCR4 locus**. A 2% TAE agarose gel was run at 90V for 20 minutes. In lane 1, 5 μL of DNA ladder was run. The following lanes contain HINDIII digest products of the Cas9NLS variants (with unbiotinylated donors) from the CXCR4 locus (5 μL in each lane). Lanes 2-5 show Cas9NLS using the CF1, Fas1, Fas2 and CF3 donors respectively. Lanes 6 and 7 are repeats of Cas9NLS Fas1 and Fas2, and lanes 8 and 9 are repeats of CF1 and CF3. Lane 10 is a positive control. Lane 11 is the unedited HEK genomic band.

**Figure S8:**
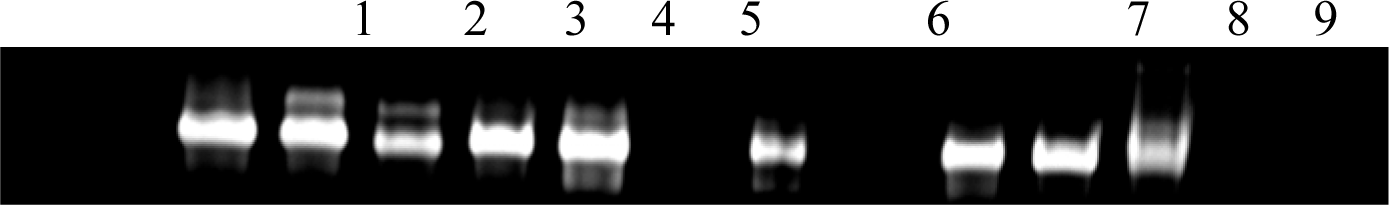
**Figure S8 Representative figure for sgRNA synthesis**. sgRNA visualised by EtBr and UV illumination. 1 FLCN1 2= FLCN 2, 3= FLCN3, 4=LBK1, 5=ACC1:1, 6=ACC1:2, 7=ACC1:3, 7=Emx1, 8=CXCR4, 9=CD81

**Figure S9:**
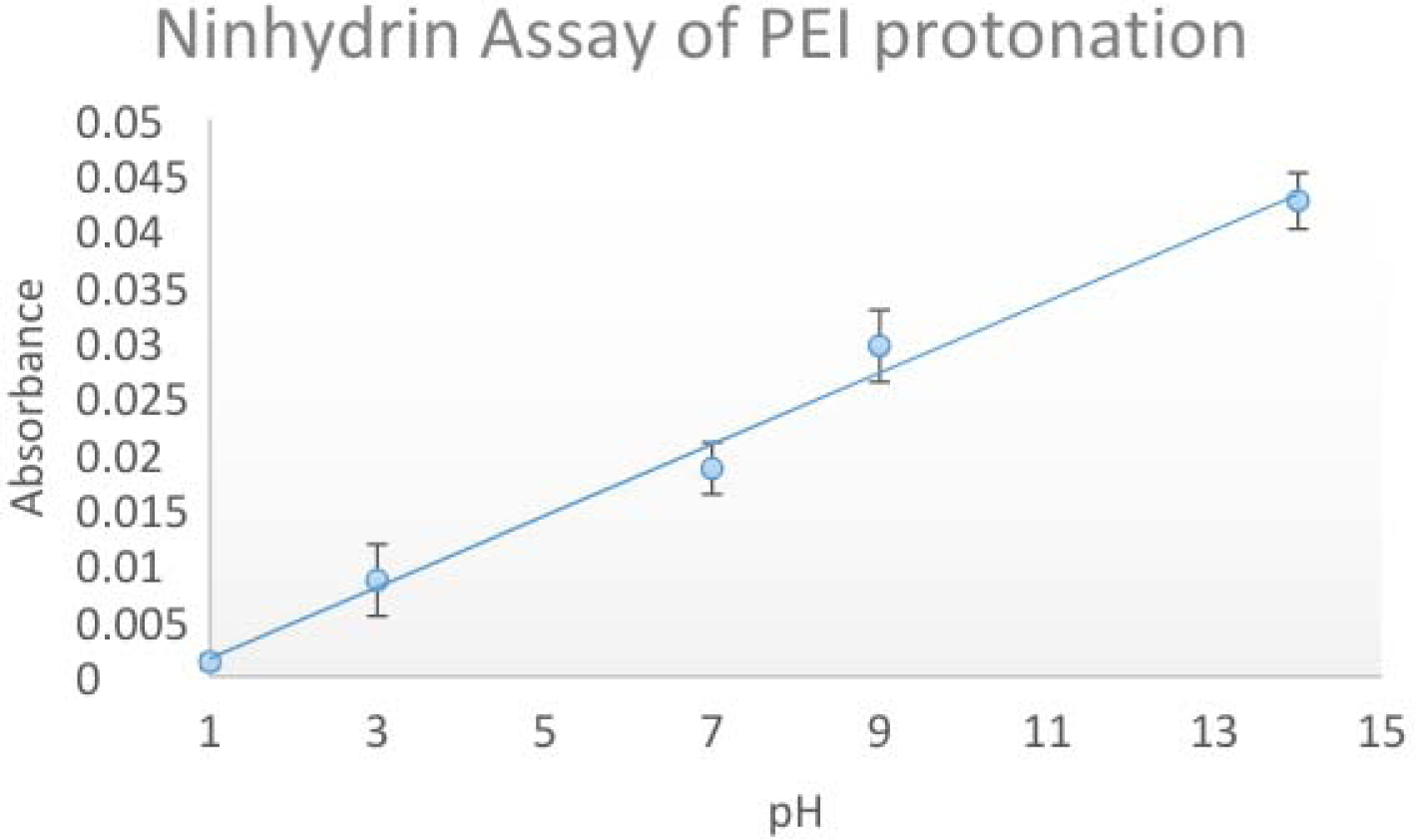
**Ninhydrin calibration curve for assessment of secondary amine protonation**

**Figure S10:**
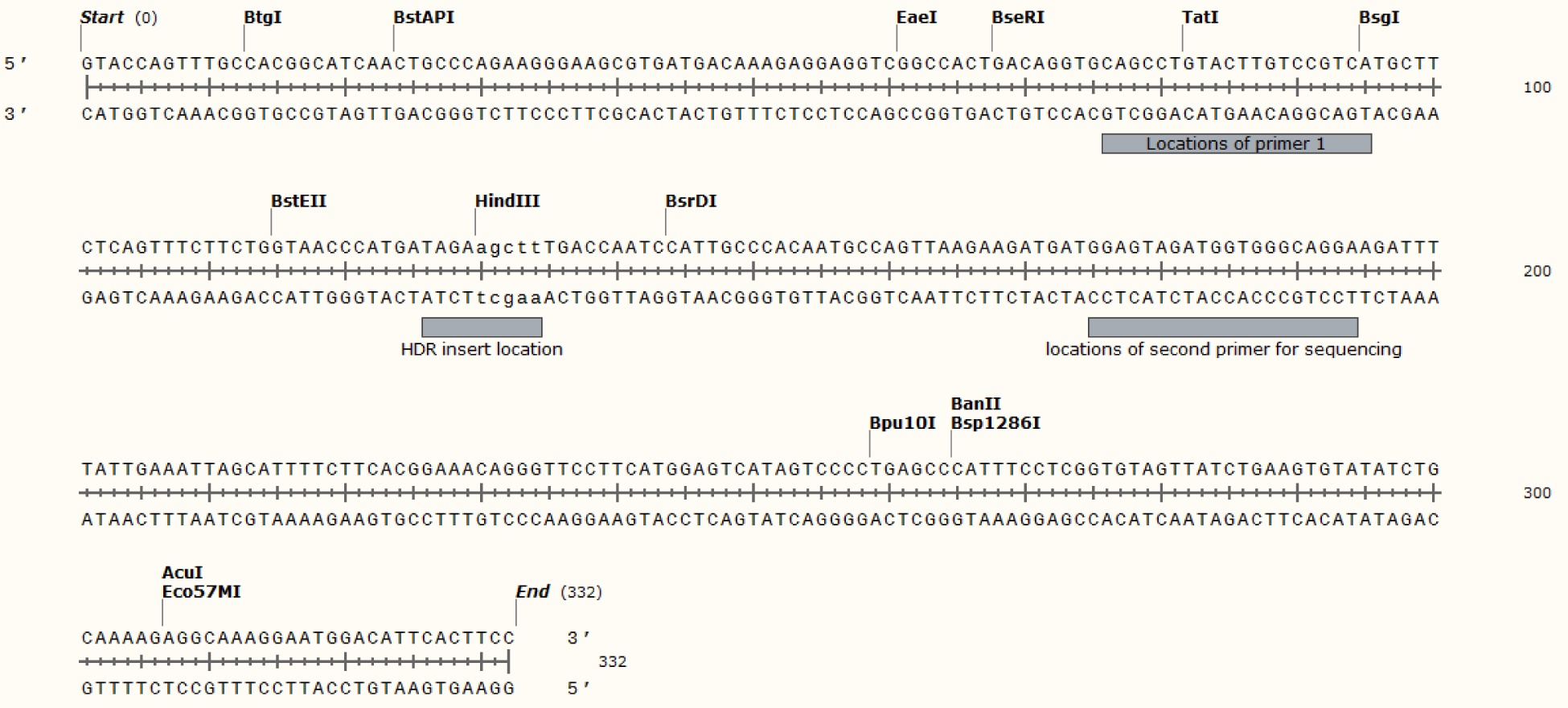
**Design of genomic primers for ion torrent sequencing**. The sequence has been edited to show the location of the proposed HindIII site.

**Figure S11:**
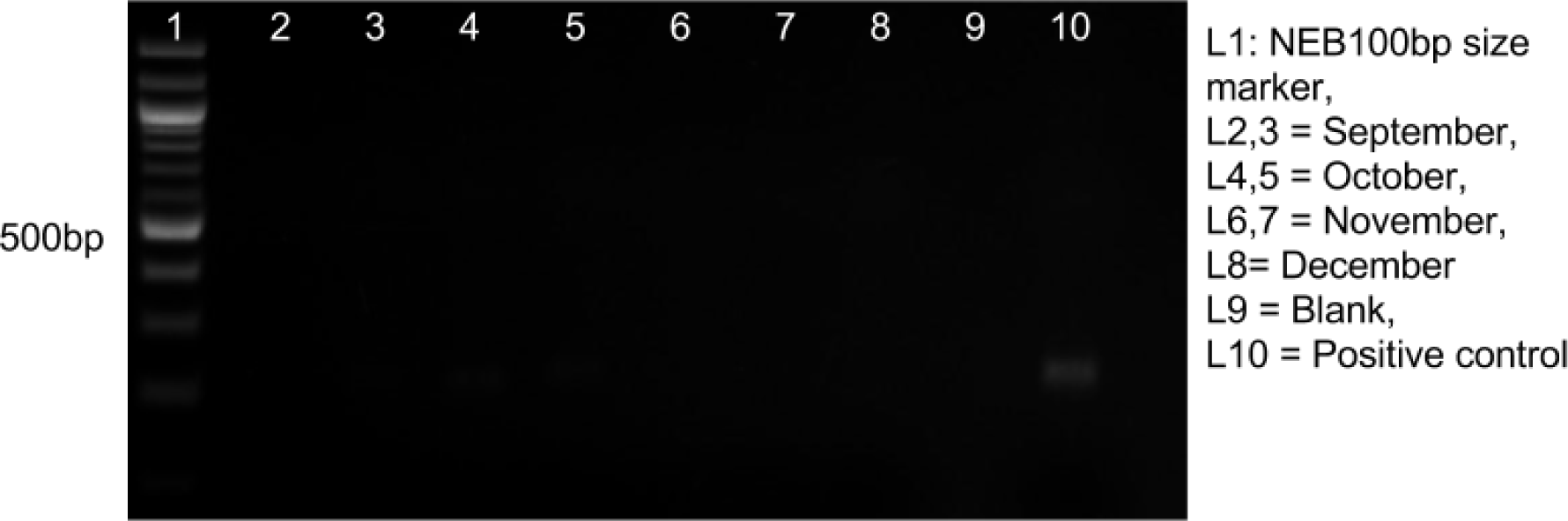
**Summary of Mycoplasma testing of Cell cultures over period of experimentation**

